# Enhancing the biological insight from limited proteolysis-coupled mass spectrometry

**DOI:** 10.1101/2024.06.12.598633

**Authors:** L. Nagel, J. Grossbach, V. Cappelletti, C. Doerig, P. Picotti, A. Beyer

## Abstract

Limited proteolysis combined with mass spectrometry (LiP-MS) facilitates probing structural changes on a proteome-wide scale *in situ*. Distinguishing the different signal contributions, such as changes in protein abundance, from protein abundance changes remains challenging. We propose a two-step approach, first removing unwanted variations from the LiP signal that are not caused by protein structural effects and subsequently inferring the effects of variables of interest on the remaining signal. Using LiP-MS data from three species we demonstrate that our framework provides a uniquely powerful approach for deconvolving LiP-MS signals and inferring protein structural changes.

## Background

The function of a protein is closely linked to its structural state, which in turn affects corresponding phenotypes^1^. Protein structural changes are important mediators of cell signaling, metabolic adaptation, molecular stress and genetic variability^2,3^. Recent technological advances have made it possible to investigate protein structural changes on a proteomic scale^4–7^. One method for simultaneously quantifying alterations in the structural state of thousands of proteins *in situ* is limited proteolysis coupled to mass spectrometry (LiP-MS). This method utilizes differences in the accessibility of protein regions to an unspecific protease, such as Proteinase K (PK)^4,8,9^. LiP-MS has shed light on a variety of biological topics including metabolic adaptation^10^, thermostability^11^, cancer-related conformational changes^12,13^, protein-protein interactions^14–16^ including proteome-specific protein-protein interactions^17^, aging-related changes in yeast^18^, *Caenorhabditis elegans*^19^ and mice^20^ as well as the use of multi-dimensional protein-structural changes as a new class of disease biomarkers^21^.

To quantify changes in the structural protein accessibility between conditions, lysates from all conditions undergo short (i.e. limited) proteinase K (PK) digestion, followed by a conventional complete trypsin digestion (LiP) (Figure 1). Differences in the protein accessibility between the conditions will reflect in distinct PK digestion patterns (Figure 1, yellow circled region) yielding altered peptide quantities. These accessibility changes may result from protein structural rearrangements or from variable shielding of surface areas, e.g., through ligand binding. Mass spectrometry (MS) is used to quantify those variations in peptide abundance on a proteome-wide scale.

**Figure 1:**
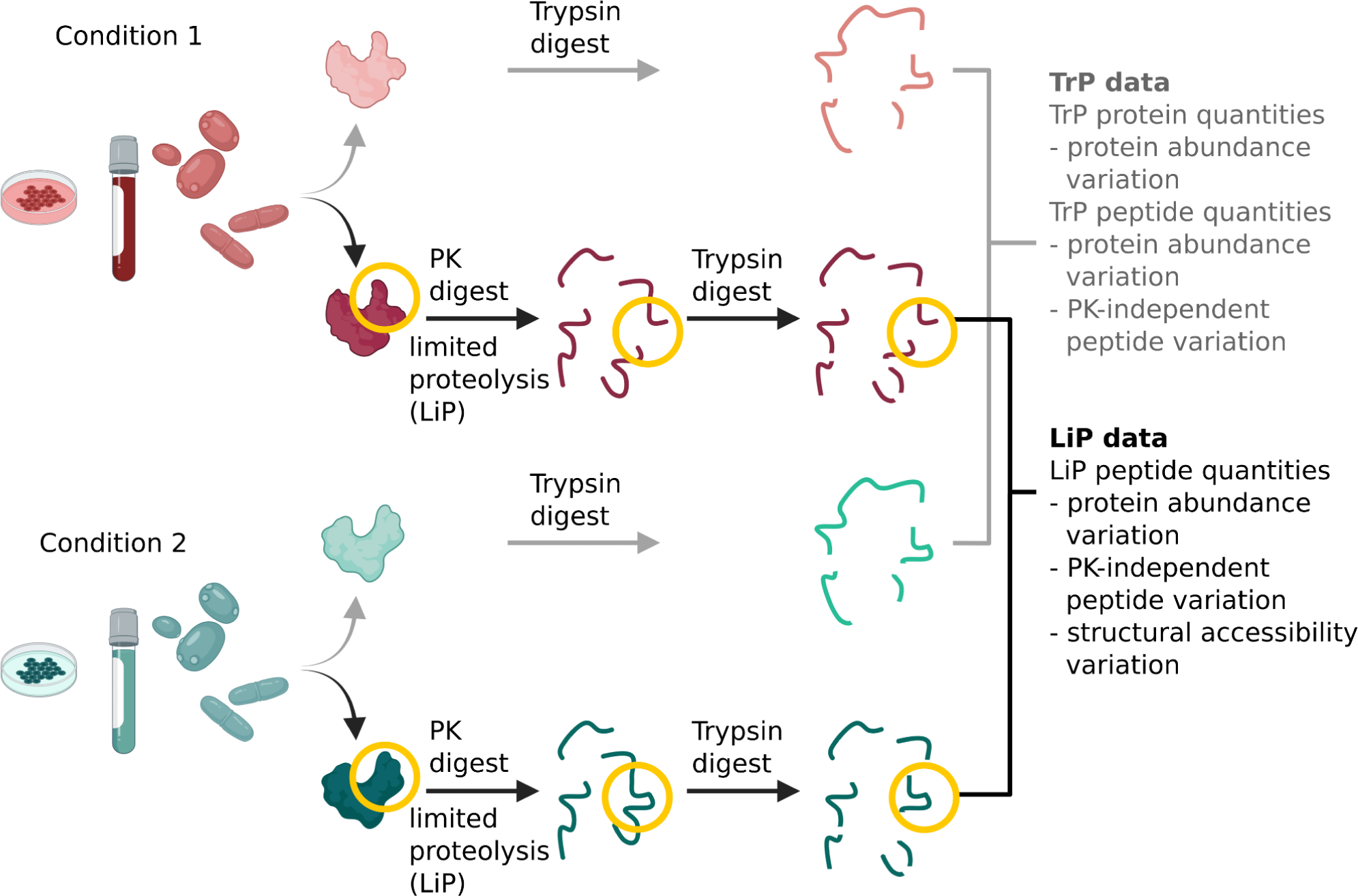
Experimental LiP-MS workflow. The proteome from samples under different conditions are extracted in nondenaturing, native conditions and digested with (a) trypsin-only (b) PK (short time, limited proteolysis) followed by trypsin. Regions with a difference in the three-dimensional conformation (yellow) show a different digestion pattern in the limited proteolysis step. Subsequently, LiP and TrP intensities are quantified via mass spectrometry. (Created with BioRender.com)

The analysis of LiP-MS data faces a number of challenges, some of which are common to molecular high-throughput data (e.g. correcting for technical variation such as batch effects, multiple hypothesis correction), while others are new and specific to LiP-MS data. First, changes in LiP peptide signals (i.e. the MS signal of a peptide subjected to the double digestion with PK and trypsin) may result from factors other than PK accessibility effects, i.e. PK-independent effects, such as variation in protein abundance, alternative splicing or post-translational modifications (PTMs). In order to account for such confounding effects, LiP-MS studies typically perform a second control digestion without limited PK digestion and only consisting of the trypsin digestion (TrP) (Figure 1). This is based on the underlying notion that PK-independent effects manifest equally in both the LiP as well as the trypsin-only digestion. Therefore, MS signal changes that are common to both data are not due to differences in the PK accessibility. Analyses schemes that appropriately combine the data from LiP-treated and TrP control samples are required to infer the signal of PK accessibility changes. Second, the PK digestion generates many half-tryptic peptides, i.e. peptides with a trypsin digestion site at one end of the peptide, while the other end was digested by PK, due to the sequence-unspecific cleavage behavior of PK. These half-tryptic peptides may carry additional information about the specific location of protein structural changes. However, their analysis is hampered by the fact that they do not exist in the trypsin-only control samples.

To address all of the LiP data specific challenges listed above, a computational approach is required that allows to distinguish the origin of variation in the LiP signal and subsequently identify the signal caused by differences in PK accessibility. As additional variation in the LiP-signal can be induced by technical effects, such as batch effects, and biological variance, analysis approaches should also enable for the removal of this unwanted variation. Existing analysis frameworks for proteomics data do not fully address all of these challenges. Approaches, typically used in the analysis of MS data that are not specifically designed for LiP-MS data^22–24^, are capable of correcting for covariates such as batch effects, but do not account for LiP-specific challenges in the data. State of the art methods designed for or previously applied to LiP-MS data do not correct for all types of unwanted variation. For example MSstatsLiP corrects LiP signals for variation caused by protein abundance changes, but it does not correct for peptide-level changes e.g. due to alternative splicing^9^. We are not aware of any tool that appropriately corrects for LiP peptide signals for all PK-independent variations.

Hence, we have developed a comprehensive method which enables the removal of unwanted variation (RUV) from LiP-MS data with different experimental setups. Our method subsequently estimated the changes in the structural accessibility between samples or conditions and is available as the R package LiPAnalyzeR. Our analysis reveals the importance of removing unwanted variation at the protein- and peptide level, thereby creating a reference framework for the analysis of LiP-MS data.

## Results

### LiPAnalyzeR - A tailored bioinformatic pipeline for inferring structural variation from LiP-MS data

The measured signal of a peptide from the LiP-MS assay (‘LiP peptide’) is affected by changes in PK accessibility and other (usually unwanted) covariates like protein abundance (Figure 3a), changes of the proteoform (e.g. alternative splicing and post-translational modifications, PTMs; Figure 3b), and technical factors such as batch effects. LiPAnalyzeR assumes that actual structural effects, reflected in changes of PK accessibility, only affect the signal of LiP peptides, while protein levels (‘TrP proteins’) and signals of peptides in the trypsin-only control measurements (‘TrP peptides’) report only on effects other than PK accessibility (Figure 3c). Therefore, signals of protein abundances (usually estimated from several peptides as single peptides are poor estimators of protein abundance^25^) and TrP peptides can be used to correct for biological signal variation of the LiP peptides that is not due to PK accessibility changes. LiPAnalyzeR removes unwanted variation (i.e. corrects for PK-unrelated covariates) by modeling the measured signal of a LiP peptide as a function of its covariates, regressing out the measured signals of those unwanted covariates, and retaining residuals containing the variation of interest. The residuals resulting from this model are utilized to estimate effects that variables of interest (e.g. different conditions) have on the structural protein accessibility in a second regression model.

Thus, LiPAnalyzeR infers structural accessibility variation in two steps (Figure 2, Figure 3): first correcting for all unwanted covariates (RUV model) and second modeling the effects of conditions of interest on the remaining signal variation (contrast model). The result of the first step is the corrected LiP signal (residual), from which the contribution of unwanted effects has been removed, and that is hence independent of protein abundance variation or variation of peptide levels not related to PK-accessibility changes. These residuals can additionally be used for further downstream analysis, such as dimensional reduction or clustering. The RUV model may also be used to correct for additional biological covariates, e.g. age, if their effect on structural accessibility variation is not of interest. The contrast model is used to estimate effect sizes, comparable to a fold change, for variables of interest and for obtaining estimates of statistical significance (p-values, see methods for details).

**Figure 2:**
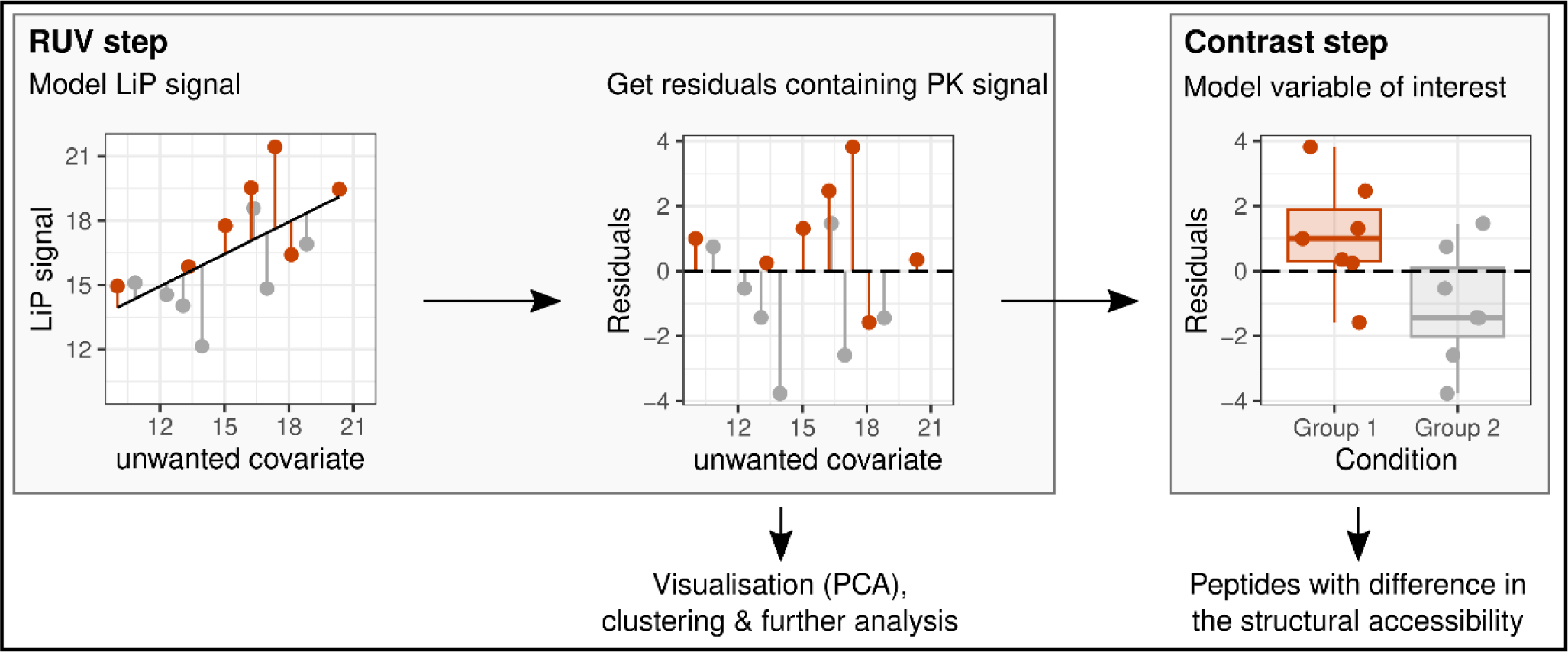
Overview LiPAnalyzeR pipeline. Covariates, which are considered a source of unwanted variation (unwanted covariates), such as protein abundances, are regressed out from the LiP signal in a RUV step (left). Residuals containing the variation of interest are estimated from the RUV models. These residuals can be used for data visualization (PCA), clustering and further downstream analysis (center). In the subsequent contrast step, the effects of the variable of interest are modeled on the residuals, identifying peptides with changes in the structural accessibility between conditions (left).

**Figure 3:**
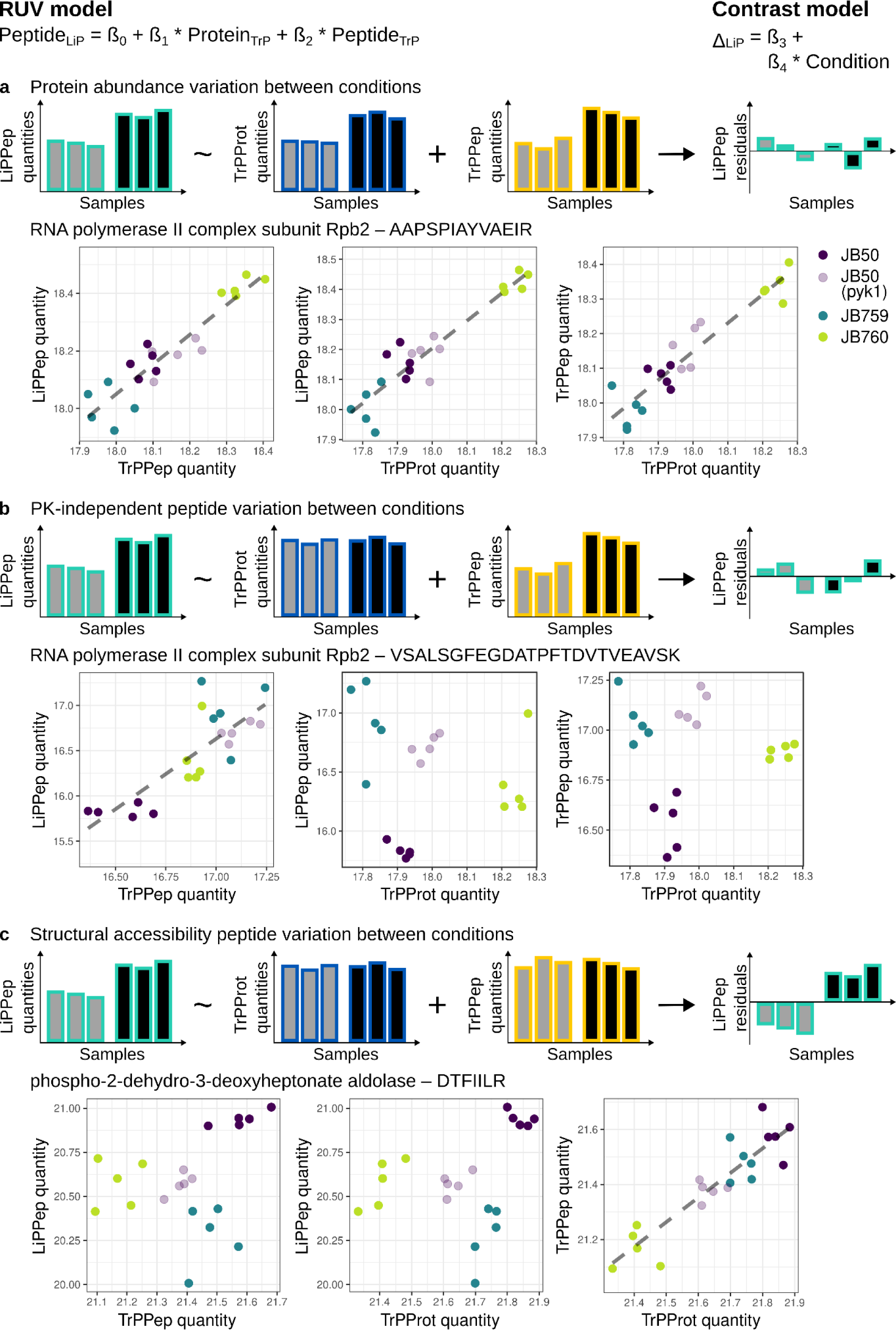
Computational approach for inferring accessibility changes between two or more conditions from LiP data. Different types of variation in LiP data are exemplified with four fission yeast strains (details see the following section and methods). **a) Protein abundance variation.** Top: Schematic overview of LiPAnalyzeR applied to a peptide containing a protein abundance effect. The RUV model removes protein abundance variation from the LiP peptides and no structural variation is inferred in the subsequent contrast model. Bottom: LiP peptide, TrP peptide and TrP protein quantities of a peptide showing a protein abundance difference between the JB50 and JB759 strain in the fission yeast data (gray dashed line visualizes linear regression). Strains are plotted in different colors (JB50: purple, PYK1 mutant: light purple, JB759: blue, JB760: green). **b) PK-independent peptide variation.** Top: Schematic overview of LiPAnalyzeR applied to a peptide with a PK-independent peptide variation. The RUV model removes PK-independent peptide variation from the LiP peptides and no structural variation is inferred in the subsequent contrast model. Bottom: LiP peptide, TrP peptide and TrP protein of a peptide showing a PK-independent peptide variation between the JB50 and JB759 strain in the fission yeast data. All colors as in a). **c) Structural accessibility variation.** Top: Schematic overview of LiPAnalyzeR applied to a peptide with a structural accessibility variation reflected only in the LiP peptides. The RUV model does not account for the structural accessibility variation, hence the signal is still reflected in the resulting residuals and detected by the contrast model. Bottom: LiP peptide, TrP peptide and TrP protein of a peptide showing variation in the structural accessibility between the JB50 and JB759 strain in the fission yeast data. All colors as in a).

### Datasets used for benchmarking LiPAnalyzeR

Multiple datasets were utilized for exploring the analysis of structural accessibility changes from LiP-MS data and benchmarking our approach implemented in LiPAnalyzeR. These datasets encompass various species and experimental setups, including multiple yeast species and human data.

We used datasets from two distantly related yeast species - fission yeast (*Schizosaccharomyces pombe*) and budding yeast (*Saccharomyces cerevisiae*). The fission yeast data consists of three different isolates (JB50, JB759, JB760) and a mutant of one of these isolates (JB50 PYK1, see methods)^26^. For each of the four strains, five biological replicates were measured in matching LiP and TrP samples. The budding yeast dataset consists of eleven biological replicates each of two well characterized isolates, measured in matching LiP and TrP samples^27^. Two technical replicates applying LiP and TrP digestion to the same sample twice were measured for every individual sample (see methods for detail).

We additionally utilized a human cerebrospinal fluid (CSF) dataset containing 52 samples from healthy individuals each measured with LiP and TrP digestion^21^. This human data set differs from the yeast data, in that the samples contain more variation due to study variables, such as age, sex or environment, and have a much more complex and varied genetic background.

### LiP peptide intensities are also affected by non-structural variation

Signals of almost all LiP peptides are positively correlated with their cognate TrP peptides and TrP proteins (Figure 4a, Extended Data Figure 1a), suggesting a strong contribution of PK-independent signals to LiP-peptide signals. This indicates the necessity to account for these PK-independent signals in the RUV step of LiPAnalyzeR. As the variation in the abundance of a protein is represented more robustly by summarizing the available levels of all TrP peptides than by a single TrP peptide^25^, TrP proteins should be used to account for protein abundance effects in the LiP data during the RUV step. We hypothesize that accounting for peptide-specific variation in addition to protein abundance variation in the RUV sep is more relevant for human samples compared to yeast samples. This is based on the fact that human proteomes are characterized by greater complexity compared to yeast proteomes, where especially alternative splicing substantially increases the diversity of proteoforms originating from the same gene. Indeed, TrP peptide signals and protein abundance were about equally correlated with LiP peptide signals in the human data, with the TrP peptides showing a slightly greater average correlation (Figure 4b). As opposed to that, in case of the fission yeast data LiP peptides signals were more closely correlated with the protein abundances, than with the matching TrP peptide signals (Extended Data Figure 1b). These PK-independent effects influence the signal of both half-tryptic and full-tryptic peptides, but are here exemplified only on full-tryptic peptides.

**Figure 4:**
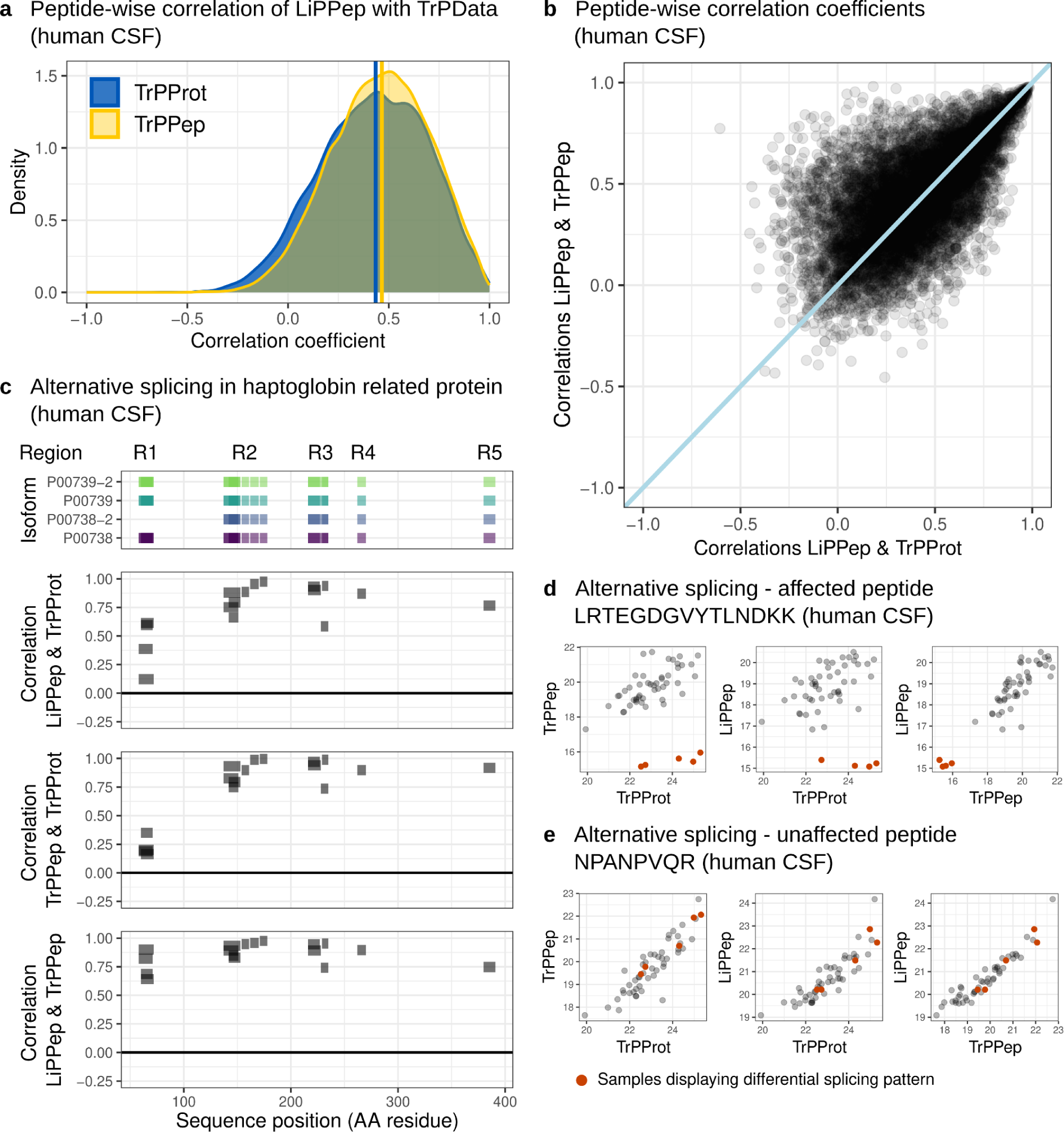
Investigation of PK-independent effects in LiP peptides, TrP peptides and TrP protein quantities. **a)** Peptide-wise Pearson correlation coefficients of LiPPep quantities with the TrPPep (yellow, median = 0.466) or TrPProt (blue, median = 0.435) quantities from the human CSF data. **b)** Correlation coefficients from a) plotted against each other. A line going through the origin with a slope of 1 is added (light blue). **c)** Alternatively spliced haptoglobin-related protein (P00739) visualized across LiP and TrP data. All peptides belonging to the main isoforms of P00739 and occuring in LiP and TrP data are visualized along the protein residues of isoforms of haptoglobin and the haptoglobin-related protein (top). Peptide-wise Pearson correlation coefficients between LiPPep & TrPProt (second from top), TrPPep & TrPProt (third from top) and LiPPep &TrPPep (bottom) are visualized along the protein residues. The protein is divided into five regions (R1-R5), where R1 is affected by alternative splicing. **d)** Example peptide from the haptoglobin-related protein affected by alternative splicing (R1). Samples with alternative splicing in R1 are red. **e)** Example peptide from the haptoglobin-related protein showing no effect by alternative splicing (R3). Samples with alternative splicing in R1 are red.

An example of PK-independent peptide variation in our human CSF data (i.e. nonrandom intensity changes in some but not all of the TrP peptides of a protein) is alternative splicing that occurs in the haptoglobin-related protein (HPR, Uniprot ID: P00739). HPR has an alternative isoform (P00739-2) and additionally a very high sequence similarity with haptoglobin (HP, Uniprot ID: P00738) and its alternative isoform (P00738-2). All peptides present in the main isoform of HPR are used to visualize a peptide-specific signal caused by alternative splicing (Figure 4c, top plot). Most HPR peptides (R2-R5) are present in all four isoforms of HPR and HP, but four peptides of Region 1 (R1, residues 58-72) are absent in isoform 2 of HP (Figure 4c, top plot). Pearson correlation coefficients were estimated for every peptide between LiP/TrP peptide and TrP protein signals across all samples (Figure 4c). Strong positive correlations were observed for all peptides from R2-R5 (mean correlations: LiP peptide to TrP protein: 0.84, TrP peptide to TrP protein: 0.90), but not for peptides from R1 (mean correlations: LiP peptide to TrP protein: 0.43, TrP peptide to TrP protein: 0.23). TrP and LiP peptides located in R1 exhibit similar deviation from the estimated protein abundances, while being highly correlated with each other (mean correlation of LiP peptide to TrP peptide R1: 0.76). This suggests that they are commonly affected by alternative splicing. Investigating the behavior of the individual samples revealed that, for samples from five donors, peptides from region R1 showed LiP and TrP peptide signals that were lower than expected based on the estimated protein abundance as opposed to peptides from the unaffected regions R2-R5; we show examples of affected (Figure 4d, red samples show an altered splicing pattern in R1) and unaffected peptides (Figure 4e). This indicates that these five donors expressed lower levels of the alternative isoform of HPR (P00739-2) compared to the other donors. Protein abundance was dominated by peptides that are common to all isoforms and hence do not reflect the alternative splicing signal. Excluding peptide-specific PK-independent signal variation from the RUV step would have falsely identified a PK-accessibility change for peptides from region R1.

These examples from fission yeast (Figure 3b) and human (Figure 4) LiP-MS datasets underline the importance of correcting for both protein- and peptide-specific – but PK-independent – variation in the LiP-MS signals. Further, the extent to which those corrections influence conclusions about specific PK effects depends on the dataset at hand.

### Strategies for removing unwanted variation from LiP-MS data

Based on biochemical principles, the PK-independent part of the LiP signal variation should be perfectly correlated with the corresponding TrP signal variation. Based on this, one should be able to correct for the PK-independent contribution by computing ratios of LiP and TrP signals (LiP/TrP ratios). In reality however, both the LiP and TrP signals are subject to measurement noise, which reduces the actual correlation between them. Simply computing LiP/TrP ratios would be suboptimal in that noise from the TrP measurements would be added to the LiP signal. The regression-based approach used by LiPAnalyzeR addresses this problem by computing the empirical association between the two measures (LiP and TrP peptide signal) for each peptide individually, thereby accounting for peptide-specific noise. To demonstrate this limitation of LiP/TrP ratios we have correlated LiP/TrP ratios with the respective TrP signals (i.e. with the matching TrP peptide and the protein abundance estimated from the TrP peptides (TrPProt); Figure 5). If the LiP/TrP ratios were corrected for the PK-independent signal variation included in both the LiP peptides and TrP measurements, the LiP/TrP ratios should show little correlation with the TrP signals. This however is not the case. Instead, the LiP/TrP ratios show strong negative correlations with the TrP peptide and TrP protein signals (Figure 5a, 5b). This bias is particularly strong for peptides where a significant structural accessibility effect is inferred based on LiP/TrP ratios, but not based on the RUV approach of LiPAnalyzeR (Extended Data Figure 1c, 1d; Supplementary text). Hence, computing ratios ‘corrects’ for variation in the TrP data that is uncorrelated with the LiP measurements, resulting in false signals.

**Figure 5:**
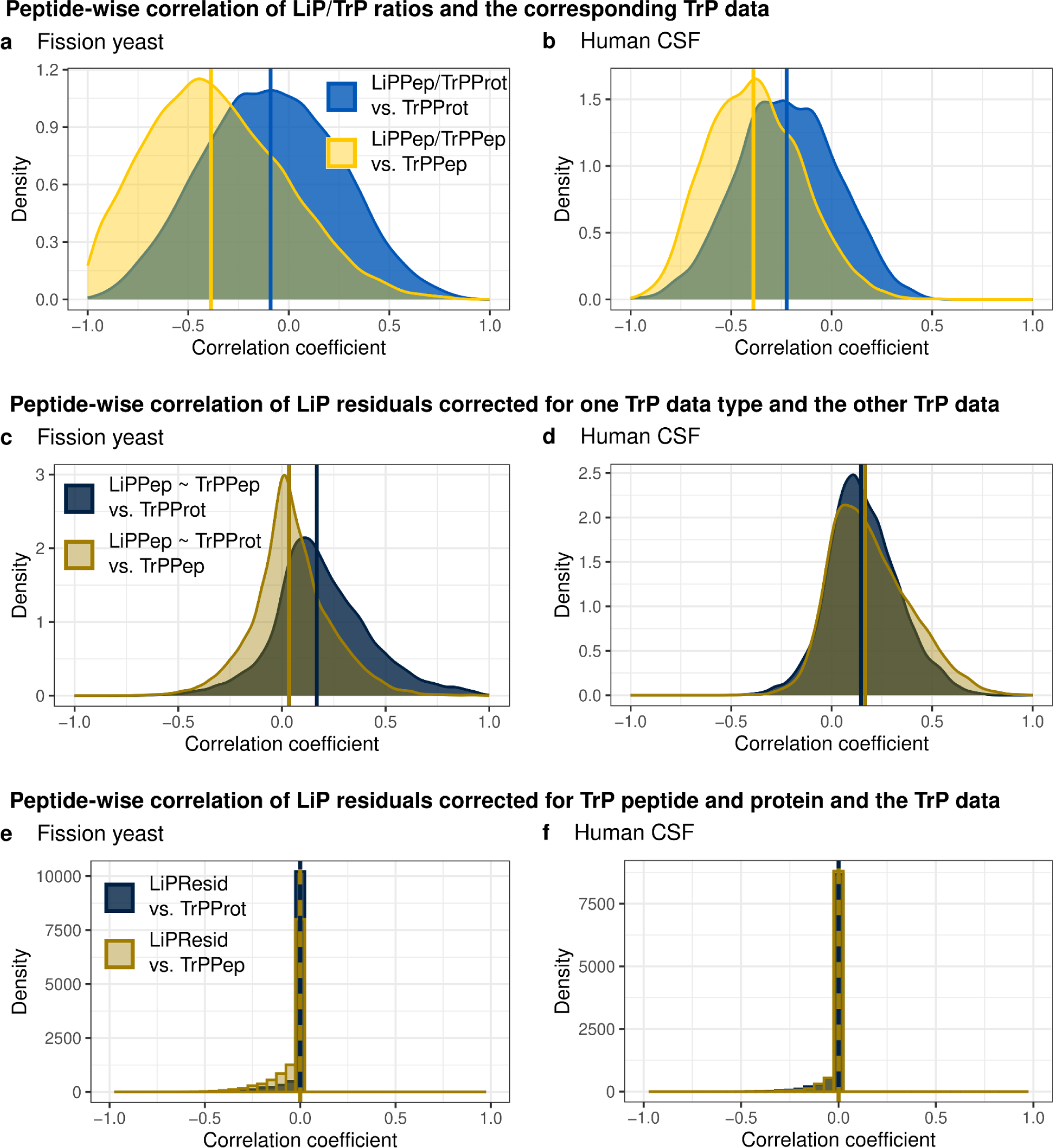
Removing unwanted PK-dependent variation from the LiP signal. **a), b)** Peptide-wise Pearson’s correlation coefficients between the ratio of LiP peptides to TrP peptides and TrP peptide quantities (yellow) as well as between the ratio of LiP peptides to TrP proteins and the TrP protein quantities (blue) in **a)** fission yeast (blue: median = −0.090, yellow: median = −0.388) and **b)** human CSF data (blue: median = −0.224, yellow: median = −0.389). **c), d)** Peptide-wise Pearson’s correlation coefficients between the residuals of the RUV step run regressing out TrP peptide or TrP protein signals from LiP peptide quantities and TrP peptide or protein quantities. Residuals estimated from models using TrP peptides as a variable are correlated against TrP protein quantities (dark blue) and residuals of those models with TrP proteins are correlated against the TrP peptide quantities (dark yellow) in both **c)** fission yeast (dark blue: median = 0.168, dark yellow: median = 0.034)) and **d)** human CSF data (dark blue: median = 0.146, dark yellow: median = 0.166). **e), f)** Residuals estimated from RUV models using both TrP peptides and proteins as variables correlated against TrP peptide (dark yellow) and TrP protein (dark blue) in **e)** fission yeast (dark blue: median = 0, dark yellow: median = 0) and **f)** human CSF data (dark blue: median = 0, dark yellow: median = 0).

The RUV model of LiPAnalyzeR corrects the signal for TrP peptide and TrP protein. In principle, the TrP peptide signal is affected by all relevant PK-independent signal variations, i.e., including variation of protein abundance and peptide effects. Therefore, it should be sufficient to correct for all PK-independent effects. However, LiPAnalyzeR includes protein abundance as an additional covariate, because single peptides are poor estimators of protein abundance^25^. Integrating the information of several peptides from the same protein reduces noise in the data. Thus, PK-independent signal variation in the LiP measurements that is mainly due to protein abundance variation is generally better corrected by using aggregate measures of protein abundance based on multiple peptides. Correcting only for TrP protein without considering TrP peptide signal would not be sufficient to remove PK-independent peptide variation from the LiP signal. To demonstrate these points, LiP peptides were corrected by running the RUV regression of LiPAnalyzeR including either TrP peptide or TrP protein signals, but not both. Pearson’s correlation coefficients were then computed between the residuals from these reduced RUV models and the respective TrP data type not used for the correction. If correcting only for TrP peptides or proteins alone is sufficient to correct for all PK-independent effects, the resulting residuals should be independent of the TrP data type not used in the RUV step. This, however, is not the case. Instead, LiP signals corrected using the reduced RUV models still retain substantial correlation with TrP protein or TrP peptide signal (Figure 5c, 5d). In the case of fission yeast (Figure 5c, median Pearson’s correlation coefficient = 0.034), correcting only for protein abundance, but not for TrP peptides, induces a smaller bias compared to the human data (Figure 5d, median Pearson’s correlation coefficient = 0.166). This is in line with the earlier observation that the human data is more strongly affected by PK-independent peptide effects such as alternative splicing.

Applying the complete RUV model - including TrP peptide and protein quantities as covariates - to the LiP peptide completely removes bias in the overall correlations of the residuals for the vast majority of peptides (Figure 5e, 5f). However, a small negative bias remains due to the non-negative constraints LiPAnalzeR applies to the coefficients estimated for the TrP peptide and protein correction (see next section). Taken together, these results demonstrate that (1) a simple ratio approach leads to artifactual signals and (2) correcting for both TrP peptide signals and protein abundance is necessary.

### Superior performance of applying a constrained RUV model prior to the contrast model

LiPAnalyzeR constrains the coefficients of the TrP peptide and protein signal to be non-negative to reduce overfitting. Negative coefficients would be biochemically implausible, since it can be expected that PK-independent effects affect the TrP and LiP data in the same direction, which is supported by the strong positive correlation of those data (Figure 3a, Extended Data Figure 1a). For instance, it would not be plausible that an increase in the abundance of a protein would result in a decrease in the signal of one of its LiP peptides. When LiP peptide signals are modeled without constraints, many of the TrP peptide and protein coefficients become negative (Figure 6a, Extended Data Figure 2a). Importantly, in the vast majority of cases, only one of the two coefficients is negative, which strongly suggests overfitting, where a too extreme positive coefficient estimated for one of the TrP data is ‘balanced’ by a negative coefficient for the other.

**Figure 6:**
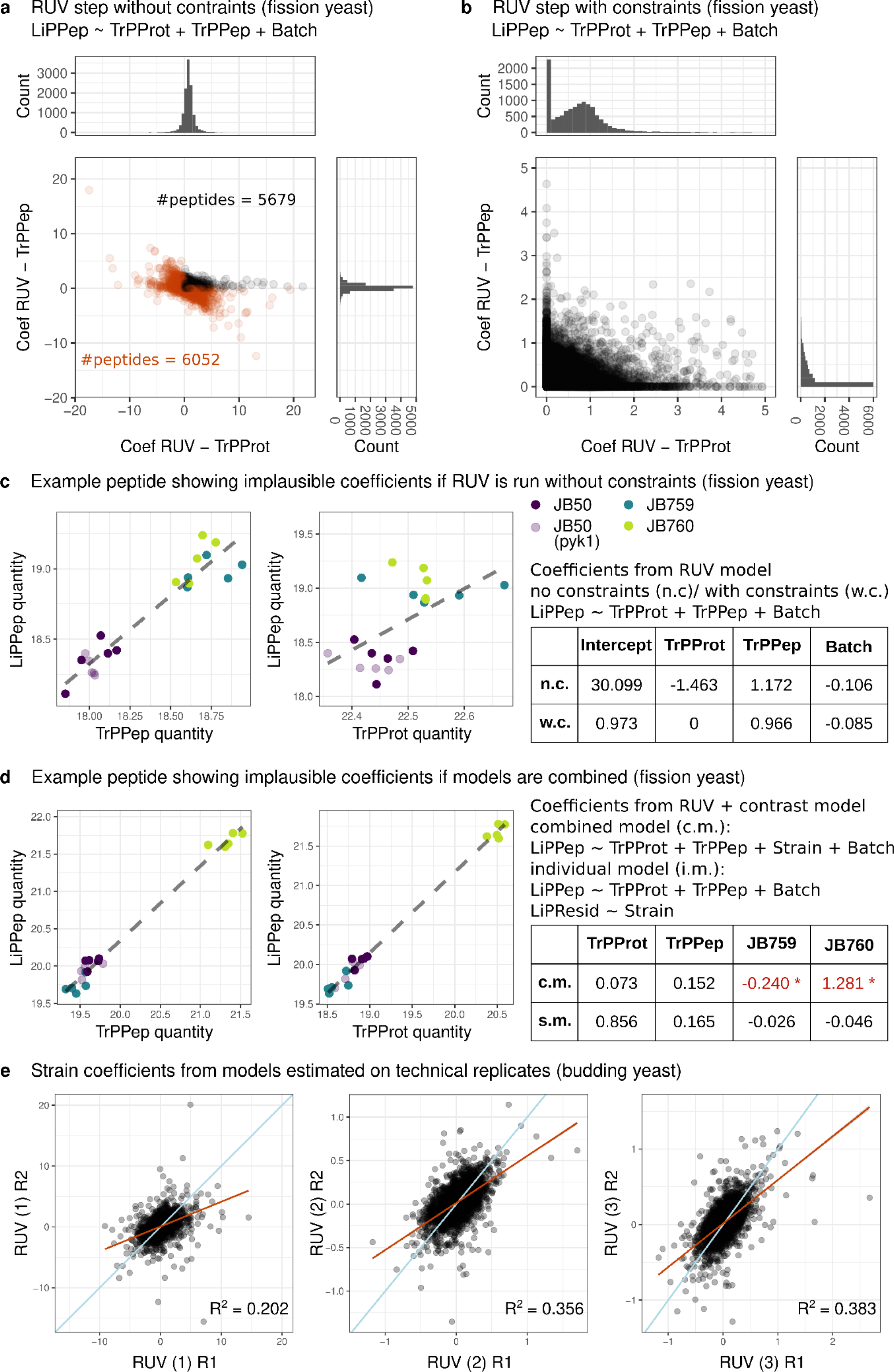
Comparison of different regression approaches applied to remove TrP signal from LiP peptide quantities. **a)** Peptide-specific coefficients for TrPPep and TrPProt from RUV models without constraints applied to fission yeast data. Peptides with at least one negative coefficient are displayed in orange. **b)** Peptide-specific coefficients for TrPPep and TrPProt from RUV models with constraints (coefficients of TrPPep and TrPProt >=0) in fission yeast data. **c)** LiP peptide quantities plotted against TrP peptide (left) and TrP protein (middle) quantities for the peptide RALIDSPCSEFPR from 60s ribosomal protein L14. Different fission yeast strains are displayed in different colors (JB50: purple, PYK1 mutant: light purple, JB759: blue, JB760: green). Coefficients from the RUV step without constraints (n.c.) and with constraints (w.c., coefficients of TrPPep and TrPProt >=0) are displayed (right), all of them have a corresponding p-value > 0.05. **d)** LiP peptide quantities plotted against TrP peptide (left) and TrP protein (middle) quantities for the peptide GLPLEAVTTIAK from dihydroxyacetone kinase Dak1 (Colors as in c)). Coefficients from a combined model consisting of RUV and contrast modeling (c.m.) and from individual models where RUV and contrast modeling is applied separately are displayed (right). Coefficients with a p-value < 0.05 are red. **e)** Peptide-wise coefficients for strain effects estimated on the technical replicates of budding yeast data using different modeling approaches: (1) combining the RUV and contrast step into one model (Equation 5, no constraints to the model), (2) running RUV without constraints and subsequently the contrast model on the resulting residuals (Equations 1-4, no constraints in model 1) and (3) running RUV with constraints and subsequently the contrast model on the resulting residuals (Equations 1-4, constraints in model 1 as described in method section). A regression line (red) and a line going through the origin with a slope of 1 is added (light blue) is shown in each plot.

In an example peptide from the fission yeast dataset for this type of overfitting, almost all variation in the LiP peptide signal could be explained by the TrP peptide signal, whereas the protein abundance shows a poor positive correlation with the LiP peptide (Figure 6c). However, an unconstrained RUV model results in a larger positive coefficient for the TrP peptide (1.172) and a negative coefficient for the protein (−1.463), which does not represent the observed behavior of the peptide and is biochemically implausible. Setting constraints to the RUV model results in a coefficient of zero for the protein abundance and estimates the correction coefficient for the TrP peptide to be very close to 1, which is biochemically plausible. Interestingly, the constrained RUV model sets one of the two coefficients at or close to zero for the vast majority of peptides (Figure 6b, Extended Data Figure 2b, 2c, 2d). Thus, for most peptides either the TrP peptide or the protein abundance is selected, but rarely both. Noticeably, more LiP peptides are TrP peptide driven in the human CSF data than in the fission yeast data (Figure 6b, Extended Data Figure 2b).

LiPAnalyzeR first removes unwanted variation (RUV step) and subsequently estimates effects for variables of interest (contrast step). In principle, it is possible to estimate the effects of unwanted covariates (such as PK-independent variation) together with the effects of interest in a single model, combining the RUV model and the contrast model. While technically feasible, this approach results in an increase in the number of false positive structural accessibility changes.

For instance, the average protein abundance often depends on the genotype^28^, hence the variable of interest (e.g. genotype) is confounded with a variable that is not of interest (e.g. protein abundance). As a consequence, LiP peptides from this protein are correlated with both genotype and protein abundance, even if the folding of the protein remains unaffected by the genotype. Combining both variables in a single model can result in modeling the protein abundance signal in the genotype (here: strain) variable, as it correlates with the LiP peptide signal. A combined model falsely assumes an effect of the genotype on the LiP peptide instead of correctly estimating the signal of protein abundance variation (Figure 6d). Applying the two-step approach first completely removes the protein abundance effect and subsequently no genotype effect is inferred in the contrast model, resulting in a more conservative approach (Figure 6d). We demonstrate that the two-step constraint regression approach chosen for LiPAnalyzeR leads to more reproducible results by systematically explored three different modeling options: (1) combining the RUV and contrast step into one model (Equation 5, no constraints to the model), (2) running RUV without constraints and subsequently the contrast model on the resulting residuals (Equations 1-4, no constraints in model 1) and (3) running RUV with constraints and subsequently the contrast model on the resulting residuals (Equations 1-4, constraints in model 1 as described in method section). We then used the consistency of strain coefficients estimated in the contrast models from different sets of replicates as a quality measure for the models: if the coefficients are more similar between replicates, the modeling approach is more robust and less prone to overfit.

These approaches were applied to both technical and biological replicates of the budding yeast dataset to compare the stability of the inferred structural variation signals (see methods for details). The strain effects estimated using the combined model were an order of magnitude larger than those resulting from the two-step approach and less consistent between replicates (Figure 6e, Extended Data Figure 3a-c). The correction coefficients for TrP peptide and protein abundance were less consistent when using a combined model (Extended Data Figure 3a-c). Further, constraining the TrP peptide and protein coefficients in the RUV model results in higher similarities between replicates compared to an unconstrained RUV model. This further supports the notion from above that an unconstrained model results in overfitting. The comparison of results from these three approaches applied to the fission yeast data led to similar outcomes (Extended Data Figure 4a-f).

Taken together, these results demonstrate that using a constrained RUV model to remove unwanted variation and applying a contrast model in a second step leads to more conservative effect estimates and reduces overfitting.

### Including half-tryptic peptides can improve coverage and sensitivity

The PK digestion of the LiP protocol produces many half-tryptic peptides. Hence, for most proteins both full-tryptic and half-tryptic peptides can be measured, resulting in a complex pattern of many peptides spanning the protein sequence (Figure 7a). For some protein regions no matching peptides are existent in the data, other regions are only covered by full-typtic or half-tryptic peptides, while some regions are covered by both full-tryptic and half-tryptic peptides. Often not only a single half-tryptic peptide matches a full-tryptic peptide, but multiple half-tryptic peptides match one or multiple (overlapping) full-tryptic peptides. For some half-tryptic peptides no matching full-tryptic peptide is present in the data. Since these PK induced half-tryptic peptides do not exist in the trypsin-only control samples, half-tryptic LiP peptides, unlike full-tryptic LiP peptides, do not have a sequence identical corresponding TrP peptide.

**Figure 7:**
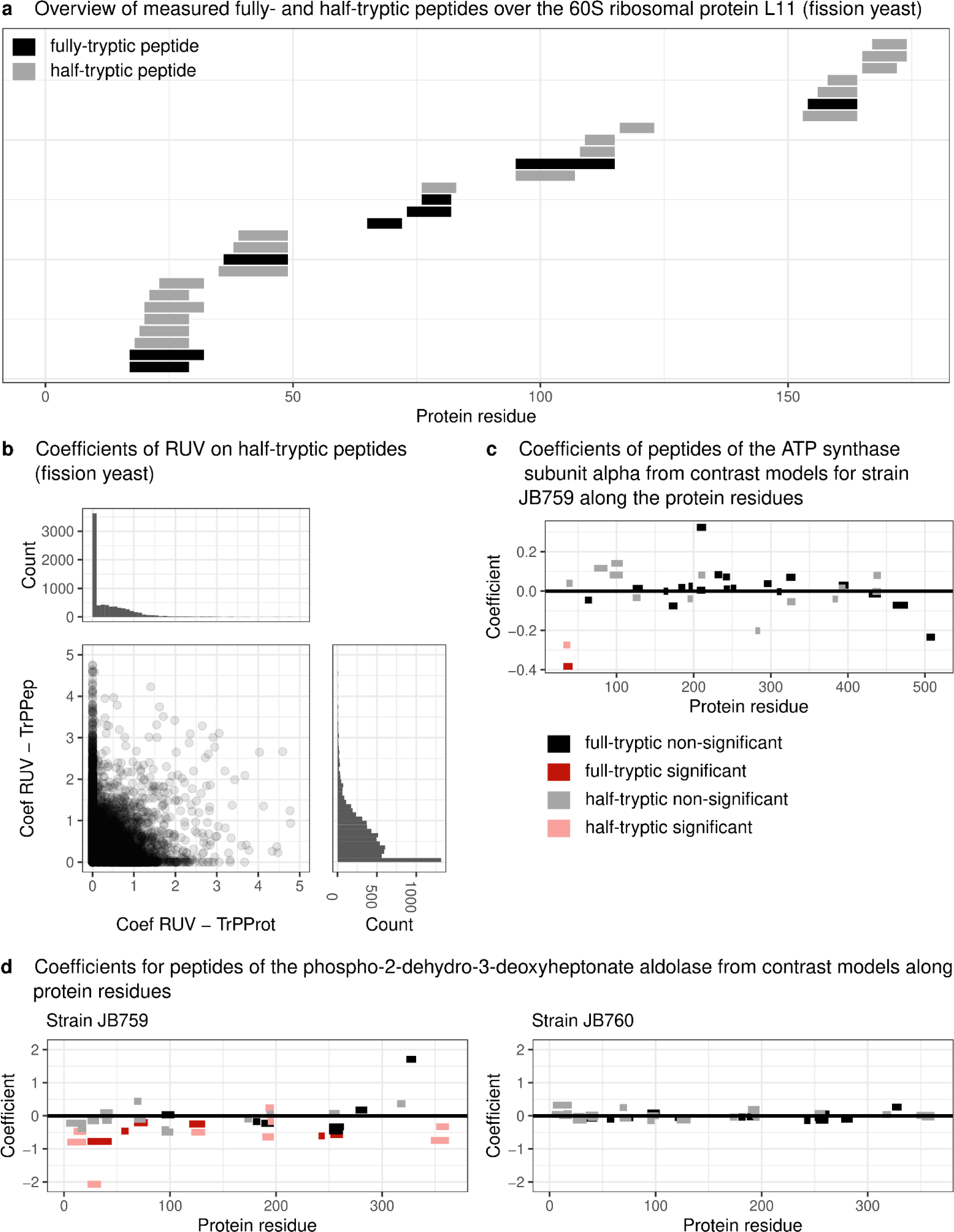
Inferring changes in the structural accessibility of half-tryptic peptides. **a)** Full-tryptic (black) and half-tryptic (gray) peptides detected in the 60S ribosomal protein L11 in fission yeast plotted along the protein residues. **b)** Peptide-specific TrPPep and TrPProt coefficients estimated by the RUV models with constraints (coefficients of TrPPep and TrPProt >=0) in fission yeast. **c)** Strain coefficients estimated by the contrast models for full-tryptic and half-tryptic peptides of the ATP synthase subunit alpha in the JB759 strain using JB50 as the reference level. RUV for the full-tryptic peptides (dark) was performed with the default LiPAnalyzeR pipeline, while half-tryptic peptides (light) were analyzed in the HT-only modus. Significant peptides (i.e., those with a protein-wise FDR corrected p-value < 0.05 estimated from the contrast models) are displayed in red. **d)** Strain coefficients estimated by the contrast models for full-tryptic and half-tryptic peptides of the phospho-2-dehydro-3-deoxyheptonate aldolase in the JB759 (left) and JB760 strain using JB50 as the reference level. RUV for the full-tryptic peptides was performed with the default LiPAnalyzeR pipeline, while half-tryptic peptides were analyzed in the HT-only mode. Colors as in c).

One strategy to deal with this ambiguity is to exclude half-tryptic peptides from the analysis, which leaves a part of the data unused. Therefore, LiPAnalyzeR also implements the following strategy to remove unwanted variation and then estimate coefficients of interest specifically for half-tryptic peptides. First, LiPAnalyzeR correlates the signal of a half-tryptic LiP peptide with all full-tryptic peptides from the TrP data from the corresponding protein(s). Subsequently, the TrP peptide with the highest positive correlation is used in the RUV model. All other aspects, such as correcting for protein abundance and applying the contrast model to the model residuals, are performed in the same way as for full-tryptic peptides. Using the most strongly correlated peptide for this correction introduces a sampling bias and may result in over-correcting the signal of the half-tryptic peptide. If many full-tryptic peptides have been measured for a protein in the TrP condition, one of them may correlate well with a half-tryptic LiP peptide by chance alone. The procedure shrinks the residuals towards zero and therefore takes a conservative approach. This can be seen by comparing the results of this approach with those of an alternative approach, where only the TrP protein and not the peptide quantities are accounted for in the RUV model. Adding the most correlated TrP peptide to the RUV model results in fewer structural changes being detected in half-tryptic peptides, although the overall directionality of the signal remains the same (Extended Data Figure 5d-f). It is also possible that PK-independent effects present in the half-tryptic peptide, such as a PTM, are not reflected in any full-tryptic peptide and therefore cannot be accounted for. Despite this limitation, half-tryptic peptides may still add useful information, especially when combined with the results from full-tryptic peptides from the same protein or protein region.

The half-tryptic LiP peptides show a higher average correlation to the matched full-tryptic peptides from the TrP data, than to the corresponding protein abundance (Extended Data Figure 5a, 5b). Thus, the TrP peptide is selected more frequently by the RUV model than the TrP protein, although the overall coefficients estimated for the TrP data by the RUV models are comparable with those estimated for the full-tryptic peptides (Figure 7b, Extended Data Figure 5c). The signal of the half-tryptic peptides can increase the confidence in structural rearrangements previously detected in the full-tryptic peptides. For example, if only a single full-tryptic peptide in a protein shows a change in the structural accessibility, a half-tryptic peptide in the same region that also shows a structural signal can corroborate the finding (Figure 7c). Since the full-tryptic peptides are individually selected by correlation, it is unlikely that the structural effects in the TrP peptides of that region are simply induced into the LiP residuals by the model. In the case of the aldolase in the fission yeast data, a strong structural effect spanning large regions of the protein can be detected in the full-tryptic peptides between JB50 and JB759, but not between JB50 and JB760 (Figure 7d). The same pattern can be observed just as strongly in the half-tryptic peptides, with there being numerous significantly changing peptides in JB759, but none in JB760. This shows how stable half-tryptic peptides can represent a structural effect that is also detected in the full-tryptic peptides.

Although the quantities of half-tryptic peptides are more variability between replicates (Extended Data Figure 5g, 5h), they can still be used to gain confidence in structural changes found by full-tryptic peptides or to detect structural variability in regions exclusively covered by half-tryptic peptides. The analysis of the full-tryptic peptides should still be run with the matching peptides from the TrP control samples in the RUV model prior to the contrast model. Half-tryptic peptides are analyzed in an independent LiPAnalyzeR run as described above.

### Working without trypsin-only control samples

Trypsin-only control measurements are essential for correcting for peptide-specific PK-independent effects. However, without such measurements it is still possible to correct for protein-level effects. Without trypsin-only control measurements, protein abundances may be estimated from the respective LiP measurements, assuming that the majority of peptides of a protein are not affected by structural changes (Extended Data Figure 6a). The correction for protein abundance variation in the RUV step is then performed using protein abundances estimated from the LiP data (LiPProt), while neglecting the correction of PK-independent peptide variation.

We first ran the RUV step on the fission yeast and human CSF data (1) with TrP peptide and protein quantities and (2) with LiP proteins only. Comparing the residuals from these two models showed a significantly higher correlation between the residuals in the fission yeast data compared to the human CSF data (Figure 8a). This was expected, since we observed PK-independent peptide variation to be more prominent in human CSF as compared to yeast samples before. We subsequently estimated strain effects with the contrast model, using the residuals estimated from RUV alternatives described above and obtained a high similarity between them (Pearson’s correlation coefficients of the strain coefficients estimated by LiPAnalyzeR: JB759 = 0.899, JB760 = 0.851, JB50 PYK1 mutant = 0.926). Therefore, using LiP proteins only, when no trypsin-only control is available, seems to be a reasonable proxy for the default RUV models where TrPPep and TrPProt are included for the majority of peptides.

**Figure 8:**
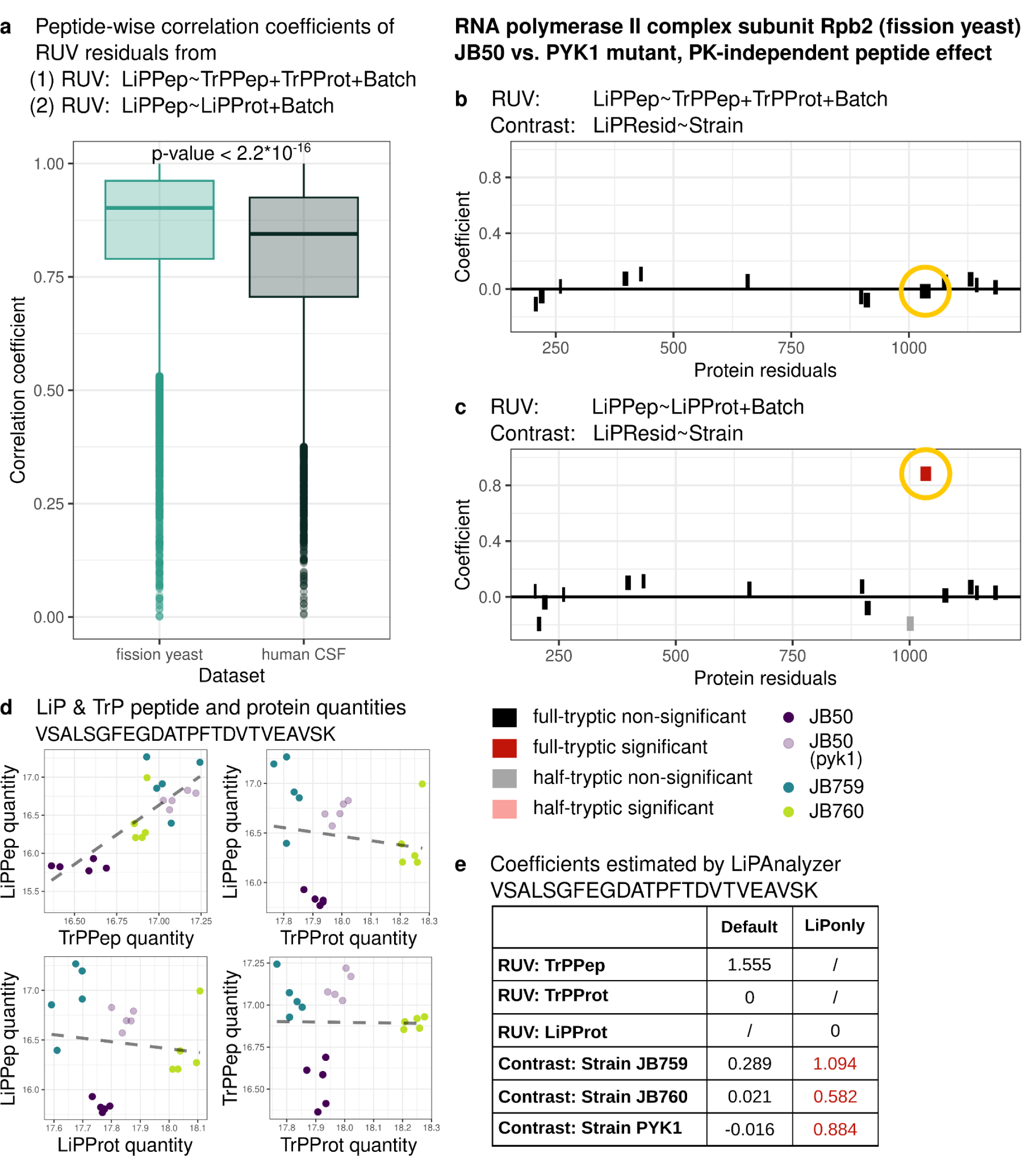
Inferring structural changes from LiP data without using the trypsin-control measurements. **a)** Peptide-wise Pearson’s correlation coefficients between the residuals from the RUV step run with (1) TrP peptide and protein quantities and (2) LiP protein quantities in fission yeast and human CSF data (Student’s t-test: p-value < 2.2*10^-16^). (Median, center; first and third quartile, lower and upper hinges; largest/smallest value no further than 1.5 × interquartile range of the hinge, whiskers; data points beyond are defined as outliers and plotted individually.) **b)** Peptide-specific coefficients of the PYK1 mutant estimated in the RUV step using the default LiPAnalyzeR analysis using JB50 as the reference strain plotted along the protein residuals. Full-tryptic peptides (dark) and half-tryptic peptides (light) are depicted, peptides with a protein-wise FDR corrected p-value < 0.05 are displayed in red. **c)** As b), but the RUV models were run in the LiP only mode, only correction for LiPProt signal. **d)** LiP and TrP peptide and protein quantities of VSALSGFEGDATPFTDVTVEAVSK plotted against each other. Colors indicate different strains (JB50: purple, PYK1 mutant: light purple, JB759: blue, JB760: green). **e)** Table with coefficients estimated by LiPAnalyzeR for the fission yeast peptide VSALSGFEGDATPFTDVTVEAVSK with the RUV step using(1) TrP peptide and protein quantities and (2) LiP protein quantities. Coefficients with a significant p-value are depicted in red.

Running the RUV model utilizing the TrP measurements of the RNA polymerase II complex subunit Rpb2 (RPB2) in the fission yeast data leads to no strain-dependent difference in structural accessibility being inferred by the subsequent contrast model (Figure 6b, 6e). On the contrary, if the RUV step is run using only the LiP protein estimates the contrast model infers a structurally accessibility variation between the strains for the peptide VSALSGFEGDATPFTDVTVEAVSK (Figure 6c, 6e). This is due to an PK-independent peptide effect which is reflected in both the LiP and TrP peptide measurements but not in protein abundance signals (Figure 6d). Therefore, not including the TrP peptide into the model prevents the RUV model from accounting for this effect, resulting in the contrast model inferring a false positive structural accessibility signal.

Utilizing LiP protein abundances instead of TrP measurements for correcting the LiP peptides is a valid option if no TrP data is at hand. However, this approach is limited by (1) the inability of correcting for PK-independent effects on peptides and (2) the presence of large scale effects, affecting a large proportion of LiP peptides, such as protein aggregation, thus affecting the estimation of LiP protein levels. How important the removal of PK-independent peptide variation is, depends on both the research question and the individual dataset/origin species.

## Discussion

LiPAnalyzeR implements a robust approach to remove unwanted variation from LiP-MS signals and subsequently identify regions in protein structures with accessibility changes. This framework consists of a RUV step that employs constrained regression, followed by a contrast model that estimates effect sizes and corresponding p-values representing PK-dependent signal changes. Our work has demonstrated that it is necessary to separate these two steps, as otherwise effects of confounding variables may be incorrectly split between a LiP effect and a confounding effect, such as protein abundance change. This ‘effect bleeding’ only occurs when two covariates are confounded, such as when batch is confounded with treatment, or protein abundance is confounded with LiP effects. Whereas the former can usually be avoided by proper experimental design, the latter is practically impossible to prevent. If a treatment affects only the abundance of a protein, it will also affect the intensity of peptides measured in the LiP assay, even in the complete absence of structural changes. An approach that jointly models the LiP signal as a function of protein abundance and treatment runs the risk of incorrectly splitting the effect between the two, even though there is no effect of the treatment on PK accessibility. This ‘effect bleeding’ can also occur when analyzing other omics data sets, such as phosphoproteomics data or when inferring alternative splicing effects from MS data. LiPAnalyzeR could additionally be applied to analysis of other types of peptide-centric structural proteomics data, such as FPOP^29^ or molecular painting data^30,31^, which are currently emerging as alternatives to LiP for the detection of structurally altered proteins in biological samples.

This work shows that TrP peptides and proteins are mostly positively correlated with the LiP peptides. Protein abundances are generally less noisy than single peptide measurements as they are estimated and averaged over multiple peptides. TrP protein abundance is therefore often a better estimator of the true LiP protein level than the observed matching TrP peptide level and is therefore often preferentially selected by the RUV model for the correction step. If, however, peptide-specific PK-independent effects (such as alternative splicing or PTMs) dominate, our modeling approach will select the matching trypsin-only peptide to correct the LiP-MS signal. Consistent with this notion, we found that TrP peptides were selected much more frequently in the comparably complex human CSF data compared to the simpler fission yeast data. Therefore, while including TrP peptides is always beneficial to minimize false positive effects, it is even more important in data where many PK-independent peptide effects are expected.

When removing both TrP peptide and protein signals from the LiP signal in the RUV step, it is crucial to prevent the corresponding coefficients from becoming negative, as negative coefficients would be biochemically implausible. Constraining coefficients within plausible ranges reduces overfitting and increases the reproducibility of the effect estimate. When the RUV step was run without constraints, over half of all peptides in budding yeast had negative coefficients. In contrast, less than one third of the peptides in the human data were affected. This is consistent with the previous observation that peptide-specific effects are less common in the yeast data. If only a protein level effect is present in the LiP data, modeling it as a function of peptide- and protein level increases the risk of overfitting. Overfitting can lead to ‘overcorrection’ when attempting to remove the influence of a confounding variable. The approach presented in this work reduces the risk of ‘overcorrection’ by imposing constraints on the RUV model. Using two variables (protein abundance and protein level) to correct for PK-independent effects is a conservative approach enabling the robust detection of PK-specific effects. Therefore, LiPAnalyzeR can also be run using either protein abundance or peptide level alone in the RUV step if this is deemed necessary for the question at hand.

The trypsin-only control experiment is used to correct for variation in the LiP-MS data that is not due to changes in the PK accessibility. Therefore, these control experiments can be omitted when no PK-independent variation in protein and peptide levels is expected, such as when mapping drug-protein binding using cell lysates^32,33^ or when structurally altered proteins are spiked into cell lysate (Extended Data Figure 7, see supplementary text). However, in all other cases, it is advantageous to have such control data. If TrP data are unavailable, protein abundances estimated from the LiP data may also be used to correct for PK-independent signal variation. The underlying notion is that changes in PK accessibility will rarely affect the entire protein. Therefore, the average signal of all peptides of a protein can serve as a proxy for the PK-independent signal of any specific peptide in the same protein. However, this approach has two disadvantages compared to having a TrP control. It is not possible to correct for peptide-specific PK-independent effects (such as splicing) because there is no independent control measurement for individual peptides. Additionally, the estimation of protein abundance from the LiP data may be flawed due to the possibility that PK digestion alters the average protein signal. In cases where proteins have only a few detected peptides, the average peptide signal may be biased by LiP effects in only one or two of these peptides. The significance of the TrP control relies on the dataset and research question at hand. If the dataset is anticipated to have more PK-independent peptide-level effects, such as splicing, it is crucial to include trypsin-control samples in the experimental setup. Our comparison of the fission yeast and human datasets confirmed this idea: omitting the TrP controls had a much greater impact on the LiP effect estimates in the human dataset than in the fission yeast data.

The RUV model produces residuals that are adjusted for confounding factors, such as protein abundance or PK-independent effects. This provides information on the structural state of the peptide. Therefore, in addition to estimating effect sizes in a contrast model, these residuals should be used for all downstream analysis focusing on LiP effects e.g., clustering of peptides based on their structural effects across conditions or visualizing structural changes.

LiPAnalyzeR is a versatile tool that can be applied beyond detecting protein structural effects. LiPAnalyzeR can also be used to detect PK-independent effects on the signal of TrP peptides by modeling TrP peptide signals as a function of protein levels. These effects might be indicative for the presence of alternative splicing or PTMs. As in the analysis of structural effects in LiP peptides, further variables such as batch membership may be included in the RUV model. LiPAnalyzeR also allows to infer differences in protein abundance between conditions using the same modeling approach as for peptide effects. These features enable the querying of a single LiP-MS dataset for a wide range of scientific questions, and associating protein structural effects with protein abundance variation and changes in proteoforms.

## Conclusions

Our work presents a unified and robust statistical framework for analyzing LiP-MS data. It addresses several challenges specific to this type of data and clearly demonstrates the need to separate effect estimation from (confounding) variable correction. This framework also allows for disentangling protein structural effects while quantifying changes in protein abundance and proteoforms, maximizing biological insight from this rich data.

## Methods

### Limited proteolysis and liquid chromatography-mass spectrometry of fission yeast Sample preparation and limited proteolysis

Five biological replicates of the Schizosaccharomyces pombe JB50, JB759, JB760 and JB50-PYK1 mutant strains were cultivated and harvested as described in Kamrad *et al.*, 2020^26^. Frozen cell pellets were resuspended in 400 μl cold LB, mixed with the same volume of acid-washed glass beads (Sigma-Aldrich), transferred to a FastPrep-24TM 5G Instrument (MP Biomedicals), and disrupted at 4°C by 8 rounds of bead-beating at 30 s with 200 s pauses between the runs. Samples were centrifuged (2 min, 1,000 g, 4°C), supernatants collected, and protein concentrations determined with the bicinchoninic acid assay (Thermo Fisher Scientific). Proteome extracts were divided into two samples: a control sample (tryptic control, TrP) undergoing only tryptic digestion to measure protein abundance changes, and a LiP sample subjected to a double-protease digestion with an unspecific protease followed by trypsin digestion to provide information on protein structural changes. Proteinase K (from Tritirachium album, Sigma Aldrich) was added to 100 µg of the LiP samples at an E:S ratio of 1:100 (w/w) and incubated for 5 min at 25°C. The same volume of water was added to 100 µg of the control samples. Digestion reactions were stopped by heating LiP samples at 99°C for 5 min, followed by the addition of sodium deoxycholate (Sigma Aldrich) to a final concentration of 5%. Control samples underwent the same procedure. Both LiP and TrP samples were then subjected to complete tryptic digestion under denaturing conditions. Peptides where reduced by incubation of samples with tris(2-carboxyethyl)phosphine (Thermo Fisher Scientific) to a final concentration of 5 mM for 30 min at 37°C. Next, the alkylation of free cysteine residues was achieved by adding iodoacetamide (Sigma Aldrich) to a final concentration of 40 mM for 30 min at 25°C in the dark. Samples were diluted with freshly prepared 0.1 M ammonium bicarbonate to a final concentration of 1% sodium deoxycholate. Samples were predigested with lysyl endopeptidase LysC (Wako Chemicals) at an enzyme/substrate ratio of 1:100. After 2 hours at 37°C, sequencing-grade porcine trypsin (Promega) was added to a final enzyme/substrate ratio of 1:100, and samples were incubated for 16 h at 37°C under shaking at 800 rpm. Protease digestion was quenched by lowering the reaction pH (< 3) The peptide mixtures were loaded onto Sep-Pak tC18 cartridges or 96 wells elution plates (Waters), desalted, and eluted with 80% acetonitrile, 0.1% formic acid. After elution from the cartridges, peptides were dried in a vacuum centrifuge, resolubilized in 0.1% formic acid, and analyzed by mass spectrometry.

#### LC-MS/MS data acquisition

Peptide digestions were analyzed on an Orbitrap Q Exactive Plus mass spectrometer (Thermo Fisher) equipped with a nanoelectrospray ion source and a nano-flow LC system (Easy-nLC 1000, Thermo Fisher). For shotgun LC-MS/MS data dependent acquisition (DDA), 1 μl peptide digestion from each biological replicate was injected at a concentration of 1 mg/ml. 1 μl of the same samples were also measured in data-independent acquisition (DIA) mode. Peptides were separated on a 40 cm x 0.75 μm i.d. column packed in-house with 1.9 μm C18 beads (Dr. Maisch Reprosil-Pur 120). For LC fractionation, buffer A was 0.1% formic acid and buffer B was 0.1% formic acid in 100% acetonitrile using a linear LC gradient from 5% to 25% or 5% to 35% acetonitrile, respectively, over 120 min and a flow rate of 300 nL/min and the column was heated to 50°C.

For DDA measurement on the Orbitrap Q Exactive Plus, MS1 scans were acquired over a mass range of 350-1500 m/z with a resolution of 70,000. The 20 most intense precursors that exceeded 1300 ion counts were selected for collision induced dissociation and the corresponding MS2 spectra were acquired at a resolution of 35000, collected for maximally 55 ms. All multiply charged ions were used to trigger MS-MS scans followed by a dynamic exclusion for 30 s. Singly charged precursor ions and ions of undefinable charged states were excluded from fragmentation.

For DIA measurements, 20 variable-width DIA isolation windows were recursively acquired. The DIA isolation setup included a 1 m/z overlap between windows, as described in (Piazza et al., 2018). DIA-MS2 spectra were acquired at a resolution of 17500 with a fixed first mass of 150 m/z and an AGC target of 1 x 106. To mimic DDA fragmentation, normalized collision energy was 25, calculated based on the doubly charged center m/z of the DIA window. Maximum injection times were automatically chosen to maximize parallelization resulting in a total duty cycle of approximately 3 s. A survey MS1 scan from 350 to 1500 m/z at a resolution of 70,000, with AGC target of 3 x 106 or 120 ms injection time was acquired in between the acquisitions of the full DIA isolation window sets.

#### Library generation and data-independent acquisition

The data were searched in Spectronaut version 15.7.220308.50606 (Biognosys). Hybrid libraries for the tryptic control and the LiP samples consisting of the corresponding DDA and DIA runs were created based on a Pulsar search using the default settings, with the exception of digestion type, which was set to “semi-specific” for the LiP samples only, and the minimal peptide length, which was set to 6. The data were searched against. The data were searched against customized fasta files^34^. The targeted data extraction was performed in Spectronaut with default settings except for the machine learning, which was set to “across experiment”, imputation which was deactivated and the data filtering, which was set to “Qvalue”. The FDR was set to 1% on peptide and protein levels. The LiP and tryptic control samples were searched separately. Results were exported using the LiPAnalyzeR_SpectroScheme format.

#### Data preparation

LiP and TrP quantities were extracted from the exported data from Spectronaut using the columns ‘*PEP.Quantity*’ for peptide and ‘*PG.Quantity*’ for protein quantities. Peptides missing a measurement in one of the samples were removed from any downstream analysis. Peptide and protein quantities were log2-transformed. Batch correction was performed on the data using the *removebatch()* function from the r-package *limma*^35^. Analysis on the data without RUV models were performed on the batch corrected data. If RUV was performed, it was run on the not batch corrected data and batch was instead added into the model as a coefficient.

### Limited proteolysis and liquid chromatography-mass spectrometry of budding yeast

#### Sample preparation, limited proteolysis and LC-MS/MS data acquisition

For this study we used 11 biological replicates of the laboratory BY4716 S. cerevisiae strain, an S288C derivative (MATα lys2Δ0) and 11 biological replicates of the wild isolate RM11-1a (MATa leu2Δ0 ura3Δ0 ho::KAN)^27^. Single colonies of the 2 strains were picked from fresh plates and inoculated in synthetic complete medium (SC,Formedium) and grown for 6 hours at 30°C under shaking at 150 rpm. The pre-cultures were inoculated into fresh SC medium cultures to a final OD600 of 0.0004 and grown overnight at 30°C under constant shaking at 150 rpm. When cultures reached OD600 = 0.8±0.1 the liquid medium was removed by 5 min centrifugation at 800 x g, RT. The pellets were washed with 20 ml of PBS 1X and centrifuged at 800 x g, RT. Cell pellets were finally resuspended in RT lysis buffer (100 mM HEPES, 1 mM MgCl2, 150 mM KCl, pH 7.5), flash-frozen and stored at −80°C. Cell lysis was performed as described above, using a FastPrep-24TM 5G Instrument (MP Biomedicals), with the following settings: speed = 5.5 m/s, time = 30 sec, 8 cycles, rest time = 200 sec. Proteome extracts were then processed as described above and peptides analyzed on an Orbitrap Q Exactive Plus mass spectrometer (Thermo Fisher), see „LC-MS/MS data acquisition“ section above.

#### Library generation and data-independent acquisition

The data were searched in Spectronaut version 15.7.220308.50606 (Biognosys). Libraries for the tryptic control and the LiP samples consisting of the DIA runs were created based on a Pulsar search using the default settings, with the exception of digestion type, which was set to “semi-specific” for the LiP samples only, and the minimal peptide length, which was set to 6. The data were searched against customized fasta files^36^. The targeted data extraction was performed in Spectronaut with default settings except for the machine learning, which was set to “across experiment”, imputation which was deactivated and the data filtering, which was set to “Qvalue”. The FDR was set to 1% on peptide and protein levels. The LiP and tryptic control samples were searched separately. Results were exported using the LiPAnalyzeR_SpectroScheme format.

#### Data preparation

LiP and TrP quantities were extracted from the exported data from Spectronaut using the columns ‘*PEP.Quantity*’ for peptide and ‘*PG.Quantity*’ for protein quantities. Technical replicates were combined taking the mean of both measurements while omitting NAs. Peptides missing a measurement in one of the samples were removed from any downstream analysis. Peptide and protein quantities were log2-transformed. Batch correction was performed on the data using the *removebatch()* function from the r-package *limma*^35^.

For comparing the results of different groups with each other, the data was split into technical and biological replicate groups. For the technical replicates, the two replicates per strain were each assigned to a different group, resulting in two technical replicates groups with eleven biological replicates of each strain in each of the groups (R1, R2). For creating the biological replicates groups (G1, G2), the summarizing technical replicates were divided by batch with G1 consisting of batch 1-5 and G2 batch 6-10.

### Limited proteolysis and liquid chromatography-mass spectrometry if human CSF

Human CSF samples were prepared and analyzed as described by Mackmull *et al.*^21^. The results from the performed Spectronaut search were exported using the LiPAnalyzeR_SpectroScheme format.

#### Data preparation

LiP and TrP quantities were extracted from the exported data from Spectronaut using the columns ‘*PEP.Quantity*’ for peptide and ‘*PG.Quantity*’ for protein quantities. Peptides missing a measurement in more than 20 samples in either the healthy or PD samples were removed from any downstream analysis. Peptide and protein quantities were log2-transformed. Batch correction was performed on the complete dataset using the *removebatch()* function from the r-package *limma*^35^. Subsequently, only the 52 healthy CSF samples were exported and used for analysis in this work.

### Limited proteolysis and liquid chromatography-mass spectrometry of spiked in α-Synuclein

Yeast samples with spiked in monomeric and fibril α-Synuclein as provided by Malinovska *et al.*^9^ (Spectronaut report LiP samples: 2019-04-29_Normalization_LiP_MSStats_Report.csv, Spectronaut report TrP samples: 2019-04-29_Normalization_Trp_MSStats_Report.csv^37^) were used.

#### Data preparation

LiP and TrP quantities were extracted from the exported data from Spectronaut using the columns ‘*PEP.Quantity*’ for peptide and ‘*PG.Quantity*’ for protein quantities. Technical replicates were combined taking the mean of both measurements while omitting NAs. Peptides missing a measurement in one of the samples were removed from any downstream analysis. Peptide and protein quantities were log2-transformed.

### Computational method of LiPAnalyzeR

The RUV step removes unwanted variation from the LiP peptide matrix *Y*_*LiP*_ = (*y*_*LiP*,*ij*_) with peptide *i* in sample *j* using a model that describes the contribution of unwanted covariables X to the measured peptide intensity:

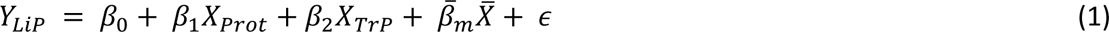

This model estimates the contribution of (whole) protein levels X_*Prot*_= (*x*_*Prot*,*ij*_), TrP peptide intensities *X_TrP_* = (*x_TrP,ij_*) and other unwanted covariates *X̅* = (*x*_*ijm*_) to the LiP signal variation of each peptide *i* independently. The matrices *X_Prot_*, *X_TrP_* and unobserved errors *ε* = *ε*_*ij*_ have the dimensionality *a* × *b*; *β*_0_, *β*_1_ and *β*_2_ are vectors of the length *a* with *β*_1_ and *β*_2_ being defined as ≥ 0; the *a* × *b* × *c* matrix *X̅* provides further variation factors for RUV and 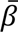_*m*_ is a matrix of *a* × *c*. Here, *a* = number of peptides, *b* = number of samples and *c* = number of further covariables for RUV.

We then estimate all the variation which can be explained by *X_Prot_*, *X_TrP_* and *X̅*:

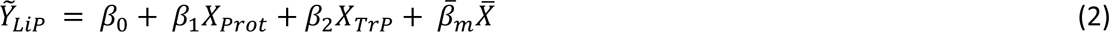

defining the *a* × *b* matrix *Ỹ_LiP_* = (*ỹ*_*LiP,ij*_) which is subsequently used to estimate the *a* × *b* residual matrix *Ŷ_LiP_* = (*ŷ*_*LiP,ij*_):

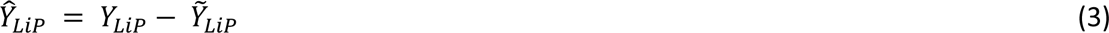

*Ŷ*_*LiP*_ contains the structural accessibility variation of interest, since all PK-dependent variation contained in trypsin-only data has been removed, as well as further variation caused by factors such as batch included in *X̅*. Accessibility variation for the covariable(s) of interest *n* can now be inferred in the contrast model

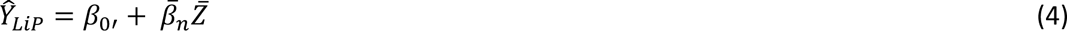

Were *Z̅* = (*z_jin_*) is a *a* × *b* × *d* matrix and 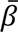*_n_* is a matrix of *a* × *d*, with *d* = number of variables of interest. 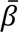_*n*_ is then evaluated via t statistics, with the degrees of freedom used in the RUV model (Equation 1) being removed from the available degrees of freedom prior to estimating the p-value. Subsequently, we correct for multiple testing effects by computing corrected p-values using the Benjamini–Hochberg approach over all peptides from the same protein (‘protein-wise FDR’).

Combining the RUV (Equation 1) and contrast (Equation 2) model into one regression (section *‘Superior performance of applying a constrained RUV model prior to the contrast model’*) results in

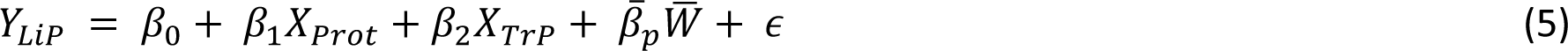

were *W̅* = (*w_jip_*) is a *a* × *b* × *e* matrix, combining *X̅* and *Y̅*, and 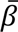_*p*_ is a matric of *a* × *e*, combining 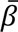_*m*_ and 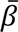_*n*_, resulting in *e* = *c* + *d*.

## Declarations

### Ethics approval and consent to participate

Not applicable

### Consent for publication

Not applicable

### Availability of data and materials

The datasets analyzed during the current study will be made available over the PRIDE database. Code for all analysis performed and the corresponding figures is provided on github: https://github.com/beyergroup/Enhancing-the-biological-insight-from-limited-proteolysis-coupled-mass-spectrometry. The R package LiPAnalyzeR can be installed from: https://github.com/beyergroup/LiPAnalyzeR.

### Authors’ contributions

LN, JG and AB conceived the project. LN conceived, designed and implemented software and analysis pipeline. VS and CD performed the LiP-MS experiments under supervision of PP. LN performed the Spectronaut searches and performed all analyses of the LiP-MS data. AB supervised the project. LN and AB wrote the initial draft of the manuscript. All authors contributed to the writing of the final manuscript.

## Acknowledgements

We thank Natalie de Souza for valuable input on the manuscript.

## Extended Data Figures

**Extended Data Figure 1:**
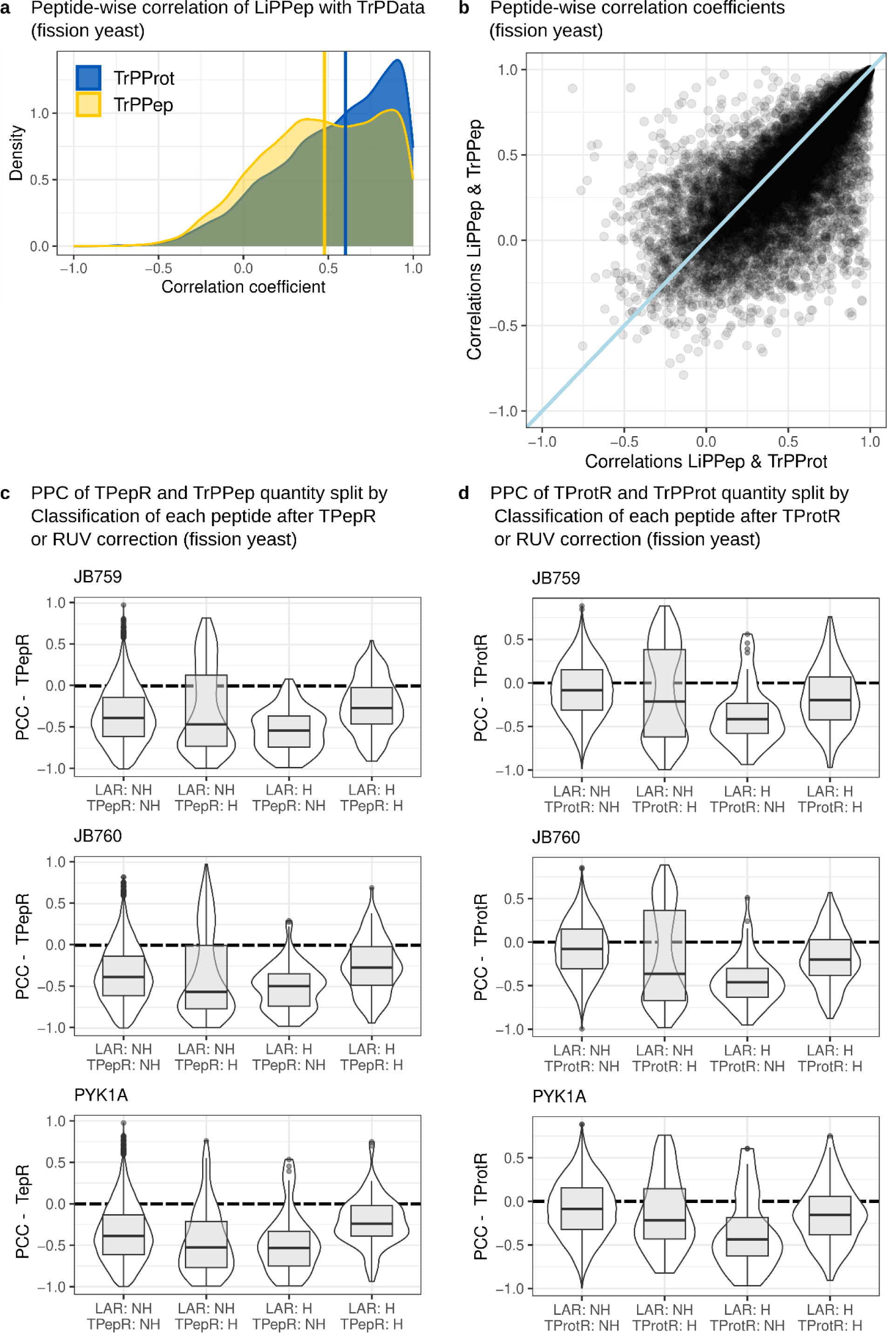
Investigation of effects across LiPPep, TrPPep and TrPProt data. **a)** Peptide-wise Pearson correlation coefficients of LiPPep quantities with the TrPPep (yellow, median = 0.478) and TrPProt (blue, median = 0.602) quantities from the fission yeast data. **b)** Correlation coefficients of a) plotted against each other for each peptide. A line going through the origin with a slope of 1 is added (light blue). **c)** Pearson’s correlation coefficient of the ratio of LiP to TrP peptides (TPepR) to TrP peptide quantities split by their classifications (NH = no hit, H = hit) using the ratio approach (TPepR) or RUV by LiPAnalyzeR (LAR) to correct for TrP quantities in the fission yeast data. Classification was performed by the contrast model of LiPAnalyzeR using JB50 as the reference strain, hence results for all other strains are shown. (For details and interpretation see supplementary text.) **d)** Same as in c but using the ratio of LiP peptides to TrP proteins (TProtR) instead. (For details and interpretation see supplementary text.)

**Extended Data Figure 2:**
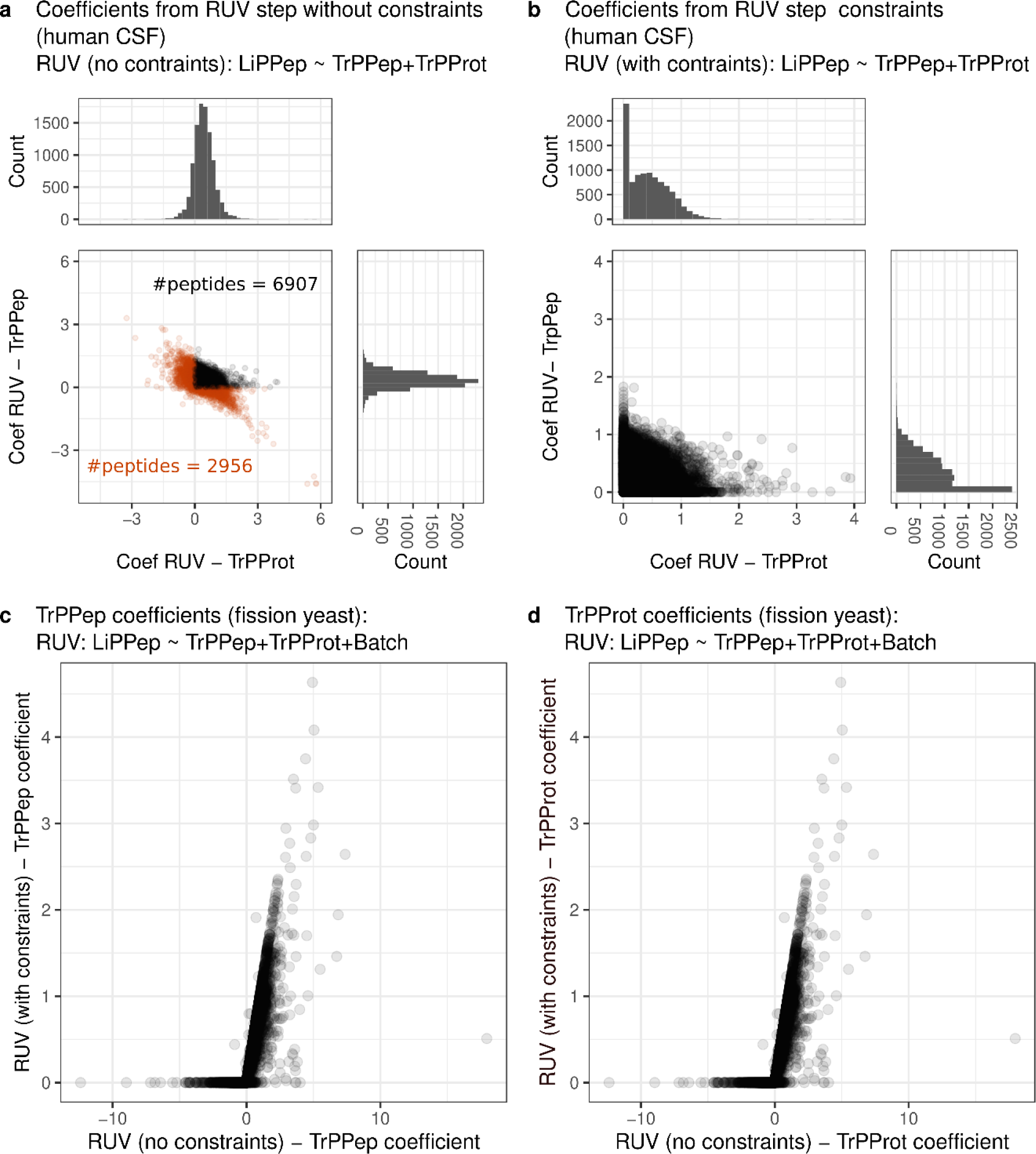
Not constraining or constraining RUV model when regressing out TrP data effects from LiP peptide quantities. **a)** Peptide-specific coefficients for TrP peptide and TrP protein quantities from the RUV step applied without setting constraints to the coefficients to the fission yeast data. Peptides with at least one negative coefficient are displayed in orange. **b)** Peptide-specific coefficients for TrPPep and TrPProt from RUV models with constraints (coefficients of TrPPep and TrPProt >=0) in human CSF. **c)** Peptide-specific coefficients for TrP peptide quantities regressed from the Lip peptide data applying the RUV step without constraints (x-axis) or with constraints (y-axis, TrP coefficients >=0) to the fission yeast data. **d)** Same as in c) but peptide-specific coefficients for TrP protein are visualized.

**Extended Data Figure 3:**
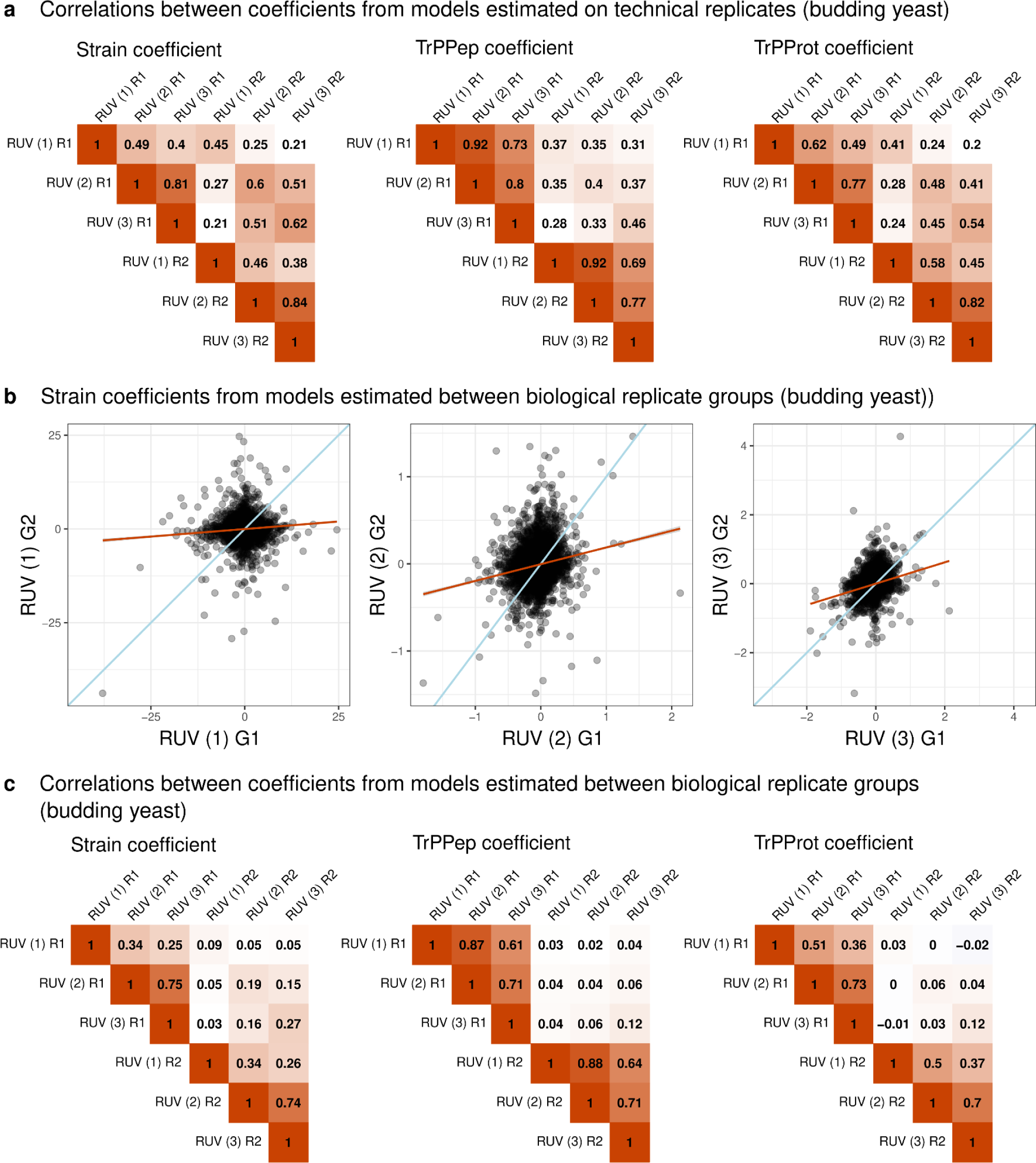
Analyzing structural accessibility variation with different RUV approaches in budding yeast replicates. **a)** Pearson’s correlation coefficients of peptide-wise coefficients estimated on the technical replicates of the budding yeast data using different modeling approaches: (1) combining the RUV and contrast step into one model (Equation 5, no constraints to the model), (2) running RUV without constraints and subsequently the contrast model on the resulting residuals (Equations 1-4, no constraints in model 1) and (3) running RUV with constraints and subsequently the contrast model on the resulting residuals (Equations 1-4, constraints in model 1 as described in method section). **b)** Peptide-wise coefficients for strain effects estimated on the biological replicate groups of budding yeast data using modeling approaches as in a). **c)** Pearson’s correlation coefficients of peptide-wise coefficients estimated on the biological replicate groups of the budding yeast data. Modeling approaches as in a).

**Extended Data Figure 4:**
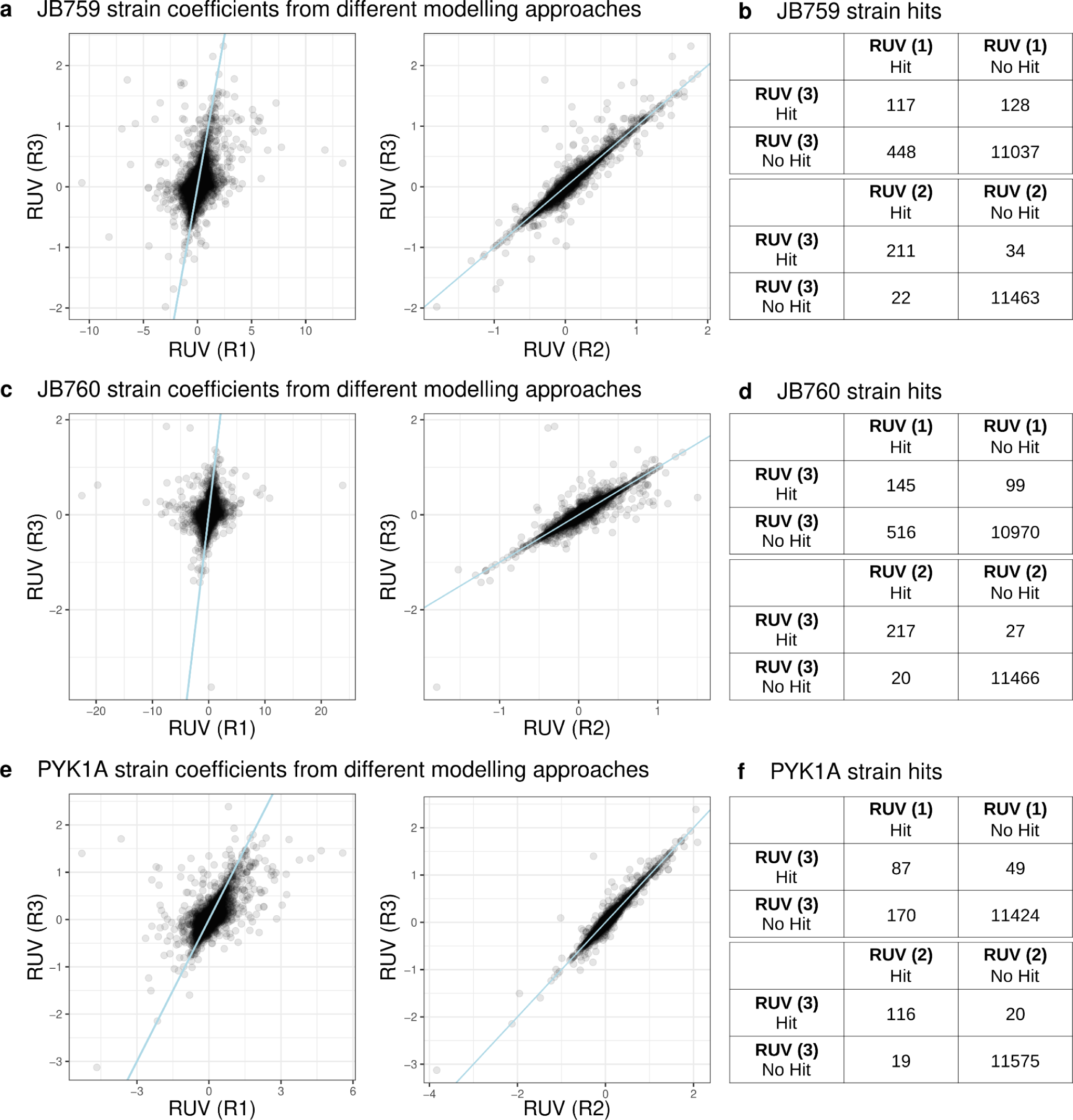
Analyzing structural accessibility variation with different RUV approaches in fission yeast replicates. **a)** Peptide-wise coefficients for the JB759 strain, with JB50 as the reference strain, using different modeling approaches: (1) combining the RUV and contrast step into one model (Equation 5, no constraints to the model), (2) running RUV without constraints and subsequently the contrast model on the resulting residuals (Equations 1-4, no constraints in model 1) and (3) running RUV with constraints and subsequently the contrast model on the resulting residuals (Equations 1-4, constraints in model 1 as described in method section). A line going through the origin with a slope of 1 is added (light blue). **b)** Number of peptides with a significant p-value for the coefficients representing the JB759 strain after protein-wise FDR when modeling the data with the same models as in a). **c)** As c) but for the JB760 strain. **d)** As c) but for the JB760 strain. **e)** As c) but for the PYK1 mutant. **f)** As c) but for the PYK1 mutant.

**Extended Data Figure 5:**
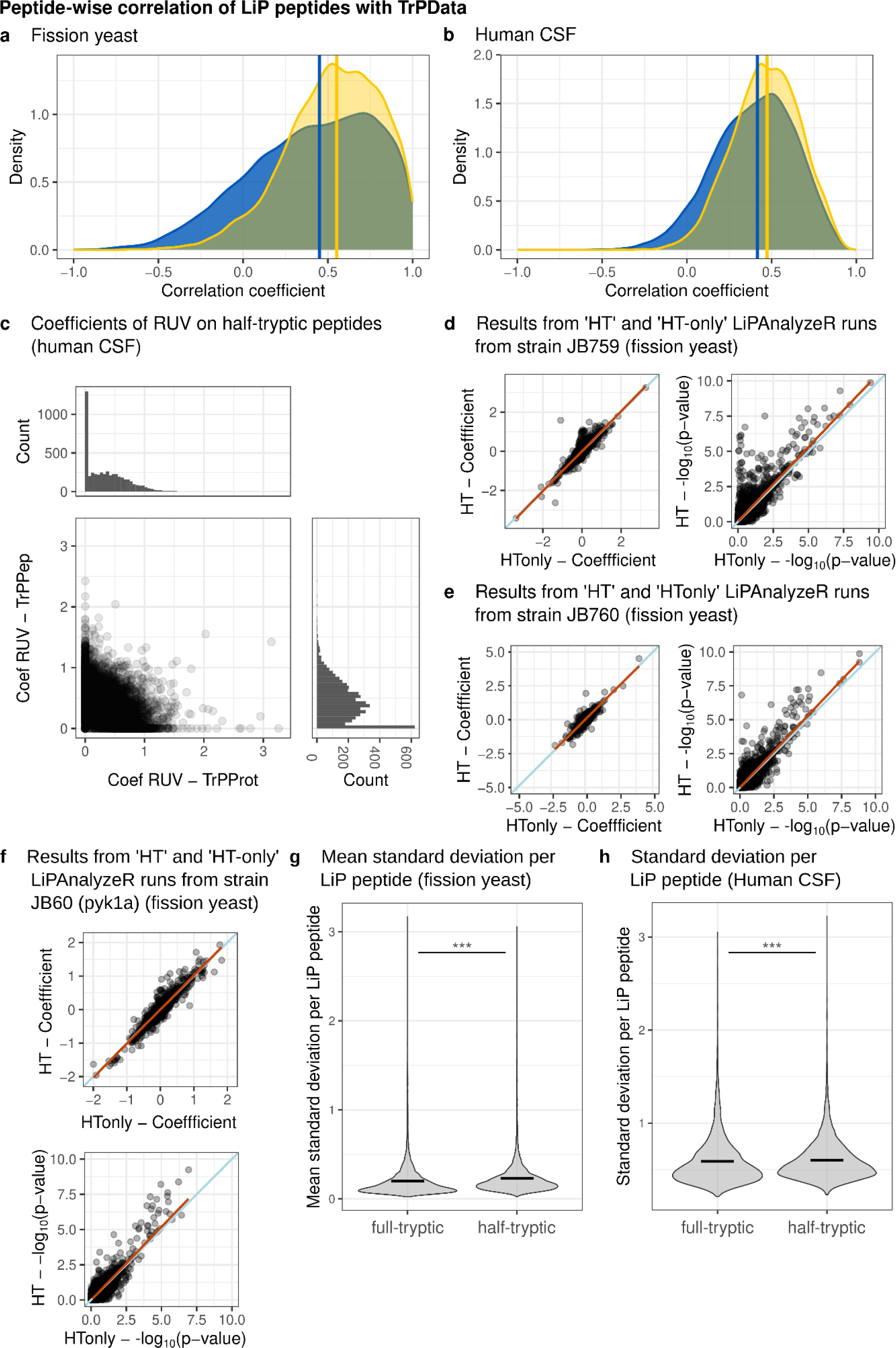
Running LiPAnalyzeR on half-tryptic peptides only. **a), b)** Peptide-wise Pearson’s correlation coefficients between the LiP peptides and and the highest correlating TrP peptide quantities (yellow) as well as between LiP peptides and the TrP protein quantities (blue) in **a)** fission yeast (yellow: median = 0.552, blue: median: 0.450) and **b)** human CSF data (yellow: median = 0.473, blue: median: 0.416). **c)** Peptide-specific coefficients for TrPPep and TrPProt from RUV models with constraints (coefficients of TrPPep and TrPProt >=0) in human CSF. **d)** Coefficients (left) and corresponding p-values (right) for half-tryptic peptides comparing strain JB759 to JB50 in the contrast model after applying the RUV model using the ‘HT’ (correcting half-tryptic LiP peptides using only the TrP protein levels) or ‘HT-only’ (correcting half-tryptic LiP peptides using the most correlating TrP peptide and the TrP protein levels). Linear regressions (orange) and lines going through the origin with a slope of 1 (light blue) are added. **e)** Same as d) but comparing the JB760 strain to JB50. **f)** Same as d) but comparing the PYK1 mutate of JB50 to JB50. **g)** Mean standard deviation of full-tryptic and half-tryptic peptides in the fission yeast data (Wilcoxon rank sum test: p-value <0.0001 (***)). The mean standard deviation is depicted as a black line. **h)** Mean standard deviation of full-tryptic and half-tryptic peptides in the human CSF data (Wilcoxon rank sum test: p-value <0.0001 (***)). The mean standard deviation is depicted as a black line.

**Extended Data Figure 6:**
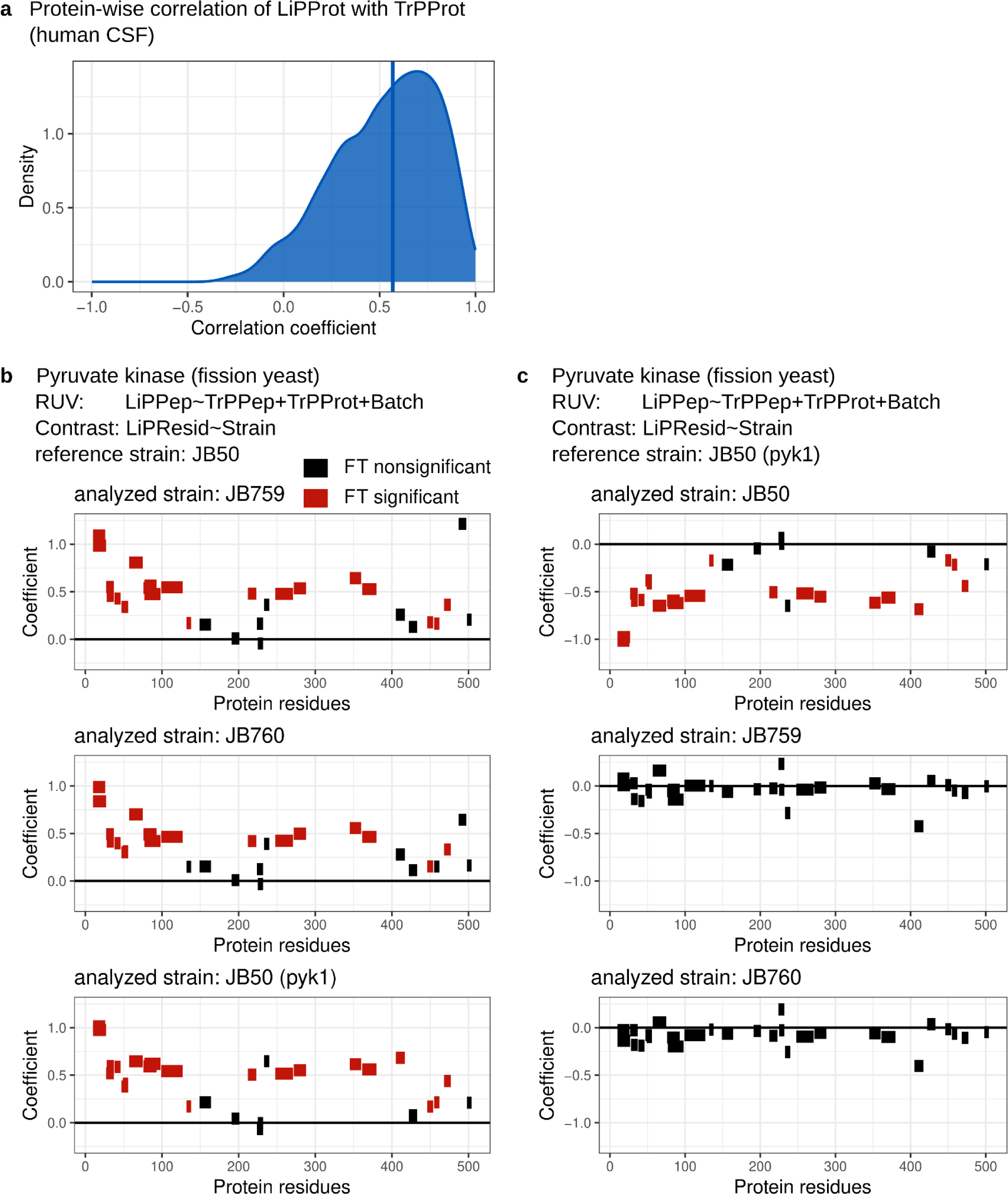
LiPAnalyzeR example results on the protein pyk1 in fission yeast. **a)** Protein-wise correlation coefficients of protein quantities estimated from TrP and LiP measurements (median = 0.569). **b)** Peptide-specific coefficients of all peptides from the pyruvate kinase estimated with LiPAnalyzeR plotted along the protein sequence for the JB759 strain (top), JB760 strain (middle) and PYK1 mutant (bottom) in the fission yeast data. LiPAnalyzeR in the default model and JB50 was set as the reference strain. Full-tryptic peptides with no significant p-value are visualized in black, if their p-value is significant they are plotted in red. (For details and interpretation see supplementary text.) **c)** As a) but, for the JB50 strain (top), JB769 strain (middle) and JB760 strain (bottom) in the fission yeast data, setting the PYK1 mutant as the reference strain. (For details and interpretation see supplementary text.)

**Extended Data Figure 7:**
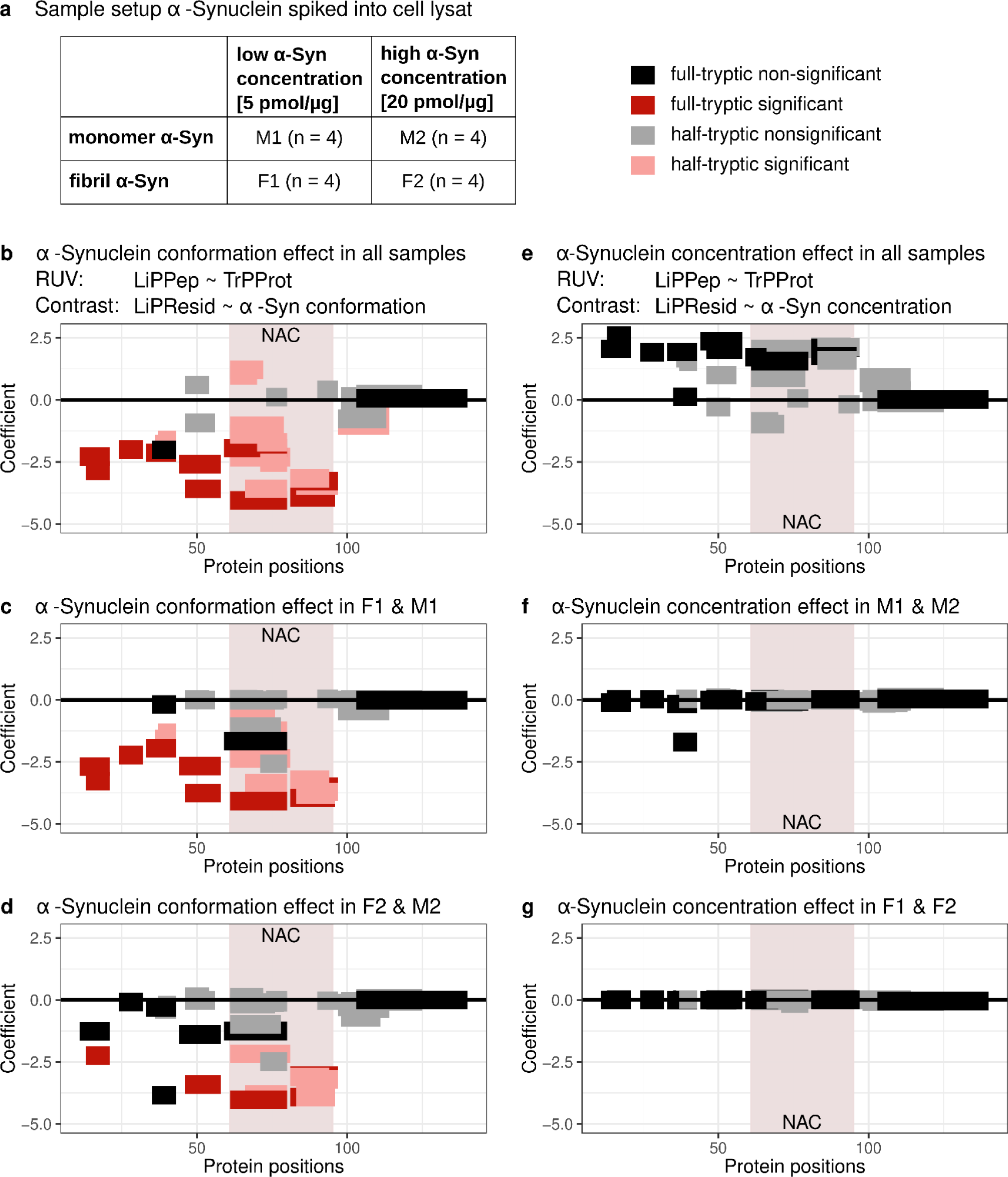
LiPAnalyzeR estimated structural effects in cell lysate spiked with monomeric and fibril α-Synuclein. **a)** Four different conditions were measured, each consisting of four biological replicates: M1: monomeric α-Synuclein in 5 pmol/μ concentration, M2: monomeric α-Synuclein in 20 pmol/μ concentration, F1: fibril α-Synuclein in 5 pmol/μ concentration, and F2: fibril α-Synuclein in 20 pmol/μ concentration. The NAC region is marked with a colored background. **b)** Peptide-specific coefficients of all α-Synuclein peptides estimated from all samples with LiPAnalyzeR plotted along the protein sequence. LiPAnalyzeR was applied on the half-tryptic and full-tryptic peptides, with the RUV model only regressing out the TrP protein abundance levels estimated from the MS data. The contrast model estimated the structural difference between monomeric and fibril α-Synuclein. Peptides with no significant p-value are visualized in black (full-tryptic) and gray (half-tryptic), peptides with a significant p-value in dark red (full-tryptic) and light red (half-tryptic). **c)** As b) but the model was only applied to the F1 and M1 samples. **d)** As b) but the model was only applied to the F2 and M2 samples. **e)** As b) but the contrast model estimated effects between the two concentration levels in which α-Synuclein was spiked into the samples. **f)** As e) but the model was only applied to the M1 and M2 samples. **g)** As e) but the model was only applied to the F1 and F2 samples. (For details and interpretation see supplementary text.)

## Supplementary text

### Ratio correction approach increases number of false classification of structural changes

To quantify the impact the use of a ratio approach to correct for TrP effects in the LiP data has on the peptides identified as being structurally altered, the ratio approach was applied to the fission yeast data. Instead of applying the RUV models, ratios of the LiP peptides to either the TrP peptides (TPepR) or TrP proteins (TProtR) were estimated and subsequently used as input for the contrast model (Equation 4), setting JB50 as the reference strain for estimating strain effects. The data was also analyzed using the standard LiPAnalyzeR pipeline (Equation 1-4). P-values estimated in the contrast model were FDR corrected protein-wise and a peptide was defined as being structurally altered if it had a corrected p-value below 0.05.

Peptides were classified based on if they were defined as structurally altered (hit) or not structurally altered (no hit) by utilizing a ratio approach (TR) before applying the contrast model or using the default LiPAnalyzeR (LAR) pipeline. This sorts the peptides into four groups for every investigated strain: 1) Hit in TR & LAR, 2) hit in TR & no hit in LAR, 3) no hit in TR & and it LAR and 4) no it in TR & LAR. Pearson’s correlation coefficients of the ratio score with the respective TrP data (Figure 4a) were plotted for each of the four groups for all strains in the TrP peptide ratio (Extended Data Figure 1c) and TrP protein ratio (Extended Data Figure 1d) version. The more negative the correlation coefficient is, the more TrP signal was transferred into the ratios and the more we expect our results to be biased by the fact that a ratio approach was applied instead of the RUV models. It can be observed that for peptides where both approaches agree (group 1 & 4) the average correlation of the ratio to the respective TrP data is much closer to zero then for peptides where there is no agreement between the approaches (group 2 & 3). Here, many more peptides have a strong negative correlation coefficient.

Hence, the more TrP signal is added into the ratio by choosing this approach, the more the final results diverge from the results we get when using our RUV model. This analysis clearly shows that using a ratio approach will increase the number of false positive and false negative peptides identified to be altered in the structural PK accessibility in further downstream analysis.

### Structural pattern of pyk1 mutation extremely stable across different strains

The pyruvate kinase (pyk1) has a single nucleotide polymorphism and associated metabolic changes in JB759, JB760 and the JB500 PYK1 mutant compared to JB50^26^. Structurally accessibility variations in peptides of pyk1 were inferred with LiPAnalyzeR using JB50 as the reference strain, revealing many peptides with altered PK accessibility (Extended Data Figure 6b, 6c). Changes in the structural accessibility of pyk1 show the same pattern over the protein residues in JB759, JB760 and the PYK1 mutant, with identical peptides in different strains being affected into the same directionality across the complete protein. If the PYK1 mutant is set as the reference strain in LiPAnalyzeR no significant structural changes are detected in pyk1 in JB759 and JB760. For JB50 structural changes are inferred in many peptides with the opposite coefficient pattern estimated for PYK1 using JB50 as the reference. The example of pyk1 demontrains the robustness with which structural accessibility changes in LiP data can be retrieved with LiPAnalyzeR.

### Alterations in the structural accessibility between monomer and fibril α-Synuclein

Monomer and fibril α-Synuclein was spiked into yeast lysate in two different concentrations (Extended Data Figure 7a, see methods for details). LiPAnalyzeR was applied to remove effects due to the different amount of spiked in α-Synuclein as well as infer differences in the structural accessibility between monomer and fibril α-Synuclein.

LiPAnalyzeR was run on all peptides of α-Synuclein measured in all samples using (1) all samples (Extended Data Figure 7b, 7e), (2) all samples with low α-Synuclein concentration (Extended Data Figure 7c), (3) all samples with high α-Synuclein concentration (Extended Data Figure 7d), (4) all samples with monomer α-Synuclein (Extended Data Figure 7f) and (5) all samples with fibril α-Synuclein (Extended Data Figure 7g). The RUV step only included the TrP protein abundance as no PK-independent peptide variations are expected in the spiked in protein. The contrast model was run modeling α-Synuclein conformation (dummy coding, reference level: monomeric α-Synuclein) for (1), (2) and (3), as well was modeling α-Synuclein concentration (dummy coding, reference level: low concentration) for (1), (4) and (5). All contrast models modeling differences in the α-Synuclein conformation estimated structural accessibility changes for many α-Synuclein peptides (Extended Data Figure 7b-d). All full-tryptic peptides mapping in the NAC region of α-Synuclein, responsible for the fibrillation of the protein^38^, display a negative coefficient, hence are more accessible in the monomers than they are in the fibrils. Contrast models modeling differences in the α-Synuclein concentration did not detect any peptides with structural accessibility changes (Extended Data Figure 7e-g), hence the RUV model removed all effects in the LiP data, based on differences in the α-Synuclein abundance.

